# Pptc7 is an essential phosphatase for promoting mammalian mitochondrial metabolism and biogenesis

**DOI:** 10.1101/426247

**Authors:** Natalie M. Niemi, Gary M. Wilson, Katherine A. Overmyer, F.-Nora Vögtle, Danielle C. Lohman, Kathryn L. Schueler, Alan D. Attie, Chris Meisinger, Joshua J. Coon, David J. Pagliarini

## Abstract

Mitochondrial proteins are replete with phosphorylation; however, the origin, abundance, and functional relevance of these modifications are largely unclear. Nonetheless, mitochondria possess multiple resident phosphatases, suggesting that protein dephosphorylation may be broadly important for mitochondrial activities. To explore this, we deleted the poorly characterized matrix phosphatase *Pptc7* from mice using CRISPR-Cas9 technology. Strikingly, *Pptc7*^−/−^ mice exhibited marked hypoketotic hypoglycemia, elevated acylcarnitines, and lactic acidosis, and died soon after birth. *Pptc7*^−/−^ tissues had significantly diminished mitochondrial size and protein content despite normal transcript levels, but consistently elevated phosphorylation on select mitochondrial proteins. These putative Pptc7 substrates include the protein translocase complex subunit Timm50, whose phosphorylation reduced import activity. We further find that phosphorylation in or near the mitochondrial targeting sequences of multiple proteins can disrupt their import rates and matrix processing. Overall, our data define Pptc7 as a protein phosphatase essential for proper mitochondrial function and biogenesis during the extrauterine transition.

---

Mitochondria are multifaceted organelles required for metabolic and signaling processes within almost every eukaryotic cell type^1^. Beyond their production of ATP through oxidative phosphorylation, mitochondria play key roles in macromolecular biosynthesis, ion homeostasis, redox signaling, and apoptotic cell death — activities that must be calibrated to changing cellular needs. Recent investigations suggest that these and other mitochondrial functions may be affected by post-translational modifications (PTMs), such as phosphorylation^2,3^ and acylation^4,5^. These PTMs are found on hundreds of mitochondrial proteins^6,7^, and can alter enzyme function^6^, complex assembly^8^, and metabolic flux^9,10^. However, other studies suggest that these mitochondrial PTMs can arise non-enzymatically^11,12^ and are often found at low stoichiometry^2,13-15^, calling into question whether and to what extent these modifications exert regulatory functions.

Although much remains to be established about the overall nature and importance of reversible phosphorylation in mitochondria, it is clear that these organelles possess a number of resident phosphatases^16-18^. For example, it has long been known that the pyruvate dehydrogenase^9^ and branched chain ketoacid dehydrogenase complexes^19^ include bound phosphatases (and kinases) that regulate their activities. However, beyond these, there exist other poorly characterized mitochondrial proteins that possess known or predicted protein phosphatase domains^16-18^, suggesting that protein dephosphorylation may be of more widespread importance in mitochondria than is currently appreciated. To begin exploring this, we recently analyzed three *S. cerevisiae* mitochondrial phosphatase deletion strains (Δ*ptc5*, Δ*ptc6*, and Δ*ptc7*)^18^ and identified phenotypes and putative substrates unique to each. In particular, we found that Δ*ptc7* exhibited a significant respiratory deficiency concomitant with elevated phosphorylation on a diverse set of proteins in the matrix where Ptc7p resides^17^. Ptc7p has also been associated with the maintenance of coenzyme Q levels^20,21^.

Here, to further explore the function of this phosphatase, we used CRISPR-Cas9 to generate a global knockout of the *PTC7* ortholog *Pptc7* in *Mus musculus.* We find that *Pptc7*^−/−^ mice are born at the expected Mendelian frequency, but exhibit severe metabolic phenotypes and die within one day of their birth. Consistent with our findings in yeast, proteomic and phosphoproteomic analyses suggest that Pptc7 influences numerous mitochondrial processes, and that its loss causes post-transcriptional downregulation of mitochondrial content and disrupts key metabolic shifts required for a successful extrauterine transition. Furthermore, our data suggest that Pptc7 influences mitochondrial protein import and processing via two mechanisms: through dephosphorylation of the mitochondrial import complex protein Timm50, and via dephosphorylation of residues within or proximal to the targeting sequences of various imported proteins. Collectively, our data argue that proper management of mitochondrial protein phosphorylation is of vital importance to mammalian metabolism and development.

## RESULTS

### Global knockout of *Pptc7* causes perinatal lethality

We generated a CRISPR-mediated global knockout of *Pptc7* in *Mus musculus.* Exons 2 (E2) and 3 (E3) were targeted simultaneously (Figure 1A), generating a founder mouse carrying indels in each region (Figure S1A). Each indel caused a frameshift that negated key catalytic residues and truncated the protein, and thus generated *bona fide* null alleles (Figure S1B). Upon breeding, the founder produced F1 progeny each carrying only one of the indels, thereby generating two independent *Pptc7*-null lines (Figure 1B). This inheritance pattern indicated that the founder mouse was compound heterozygous (null) for *Pptc7*, yet this mouse showed no overt morphological or metabolic abnormalities for ~18 months (data not shown).

**Figure 1:**
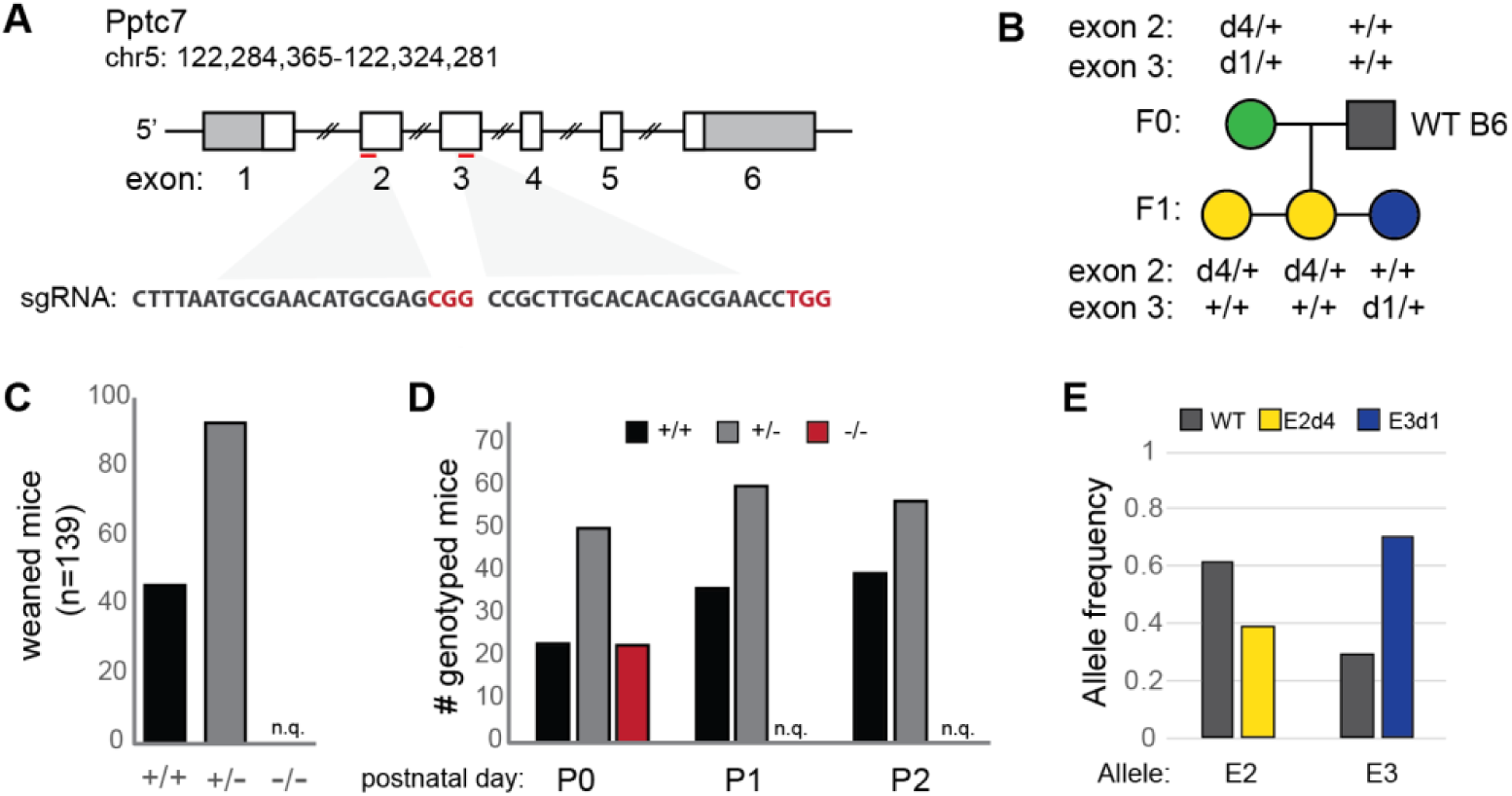
Global, CRISPR-mediated knockout of *Pptc7* causes perinatal lethality. A. Targeting strategy for the *Pptc7* locus in *Mus musculus* is shown; two sgRNAs were designed to exons 2 and 3 and injected simultaneously into 1 cell zygotes. B. The founder mouse was bred to a wild type C57B6/J mouse, generating F1 pups containing deletions (labeled dX where X is number of base pairs deleted) in exon 2 (yellow) or exon 3 (blue). This segregation pattern indicates the founder mouse is compound heterozygous (null) for *Pptc7.* C. No *Pptc7* knockout pups were found at weaning (n=130). D. *Pptc7* knockout pups are born at Mendelian frequencies, but no live pups were found at postnatal day 1 (P1) or day 2 (P2) (reported as n.q. – not quantified). E. Allele frequency in skeletal muscle of the F0 founder mouse, demonstrating mosaicism.

We interbred *Pptc7*^+/−^ mice from both lines and genotyped their progeny, but found no *Pptc7*^−/−^ (KO) mice in the weaned pups (n=130). This observation suggests that CRISPR-mediated loss of *Pptc7* via either indel causes lethality that is not due to off-target effects (Figures 1C, S1C-E). We found that pups of all genotypes are born at Mendelian frequencies (Figure 1D, S1F-H), but fail to survive after birth (Figure 1D), indicating that KO pups develop appropriately *in utero* but cannot survive the perinatal transition.

The full penetrance of perinatal lethality in the *Pptc7* knockout pups led us to investigate how the founder mouse, who was compound heterozygous for *Pptc7*, survived. CRISPR-generated mice are often mosaic^22^, suggesting that the founder may have expressed a non-Mendelian ratio of wild type and null alleles in its somatic tissues. To test this, we performed next generation sequencing (Figure SI) and found skewed allele frequencies in all tissues examined from the founder (Figure S1J-P), while F1 progeny had expected allele frequencies (Figure S1Q-T). Notably, the founder had a substantial percentage of wild type alleles (Figure 1E), which likely contributed to survival. Collectively, these data demonstrate that *Pptc7* expression is required for survival of the extrauterine transition in mice, and that CRISPR-Cas9 associated mosaicism serendipitously enabled survival of the founder mouse and germline transmission of knockout alleles for an essential gene.

### Pptc7-null mice have defects associated with inborn errors of metabolism

During birth, mammals transition from a primarily glycolytic metabolism *in utero* to a reliance on lipid- and protein-rich milk as a nutrient source^23,24^. As such, deficiency in metabolic processes including glycogen mobilization^25^, fatty acid oxidation^26^, and ketone body utilization^27^ can lead to perinatal lethality. We profiled these and other metabolic phenotypes to assess their potential contribution to the early *Pptc7*^−/−^ lethality. *Pptc7* KO pups (Figure 2A, Figure S2A-C) and E14.5 embryos (Figure S2D) weighed significantly less than their WT counterparts. *Pptc7* KO pups were also hypoglycemic (Figure 2B), with a median blood glucose of 41 mg/dl relative to 71 mg/dl for WT pups. Insulin levels were unchanged in KO relative to WT pups (Figure 2C), suggesting that their hypoglycemia may alternatively have arisen through impaired gluconeogenesis and/or increased glycolytic flux. Consistently, KO pups displayed lactic acidosis (Figure 2D), which is commonly associated with mitochondrial dysfunction^28^. Finally, KO mice were hypoketotic, with a ~3.5-fold decrease in serum ketones relative to their WT littermates (Figure 2E), and analysis of matched glucose and ketone levels across mice demonstrates hypoketotic hypoglycemia (Figure S2E).

**Figure 2:**
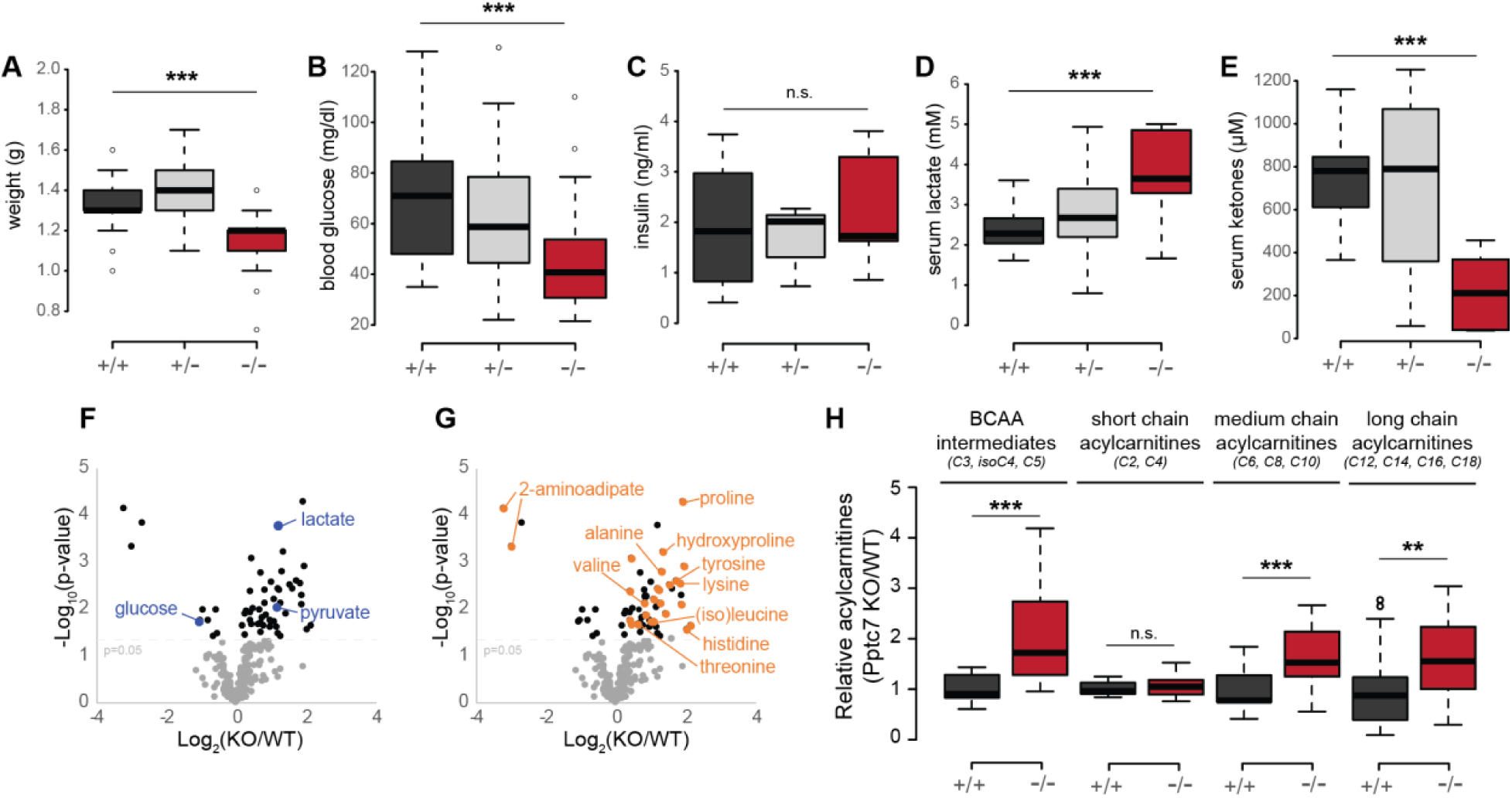
*Pptc7*-null mice have defects associated with inborn errors of metabolism. *Pptc7* KO perinatal pups (P0, red, n=46) weigh significantly less than wild type (dark gray, n=47) and heterozygous (light gray, n=102) littermates. B. *Pptc7* KO pups (n=28) are hypoglycemic relative to WT (n=41) and heterozygous (n=82) littermates C. *Pptc7* KO pups (n=10) show no difference in circulating insulin relative to WT (n=8) and heterozygous (n=5) littermates. D. *Pptc7* KO pups (n=7) have elevated serum lactate relative to WT (n=9) and heterozygous (n=22) littermates. E. *Pptc7* KO pups (n=6) have lower concentrations of serum ketones relative to WT (n=9) and heterozygous (n=13) pups. F. Metabolomics from liver tissues reveal decreased glucose and increased lactate and pyruvate in KO (n=5) samples relative to WT (n=5). G. Metabolomics from liver tissue (same data as shown in 2F) reveals numerous differences in amino acids and their intermediates in KO samples relative to WT. H. Acylcarnitine analysis of liver tissues from KO and WT. For each plot, n=7-8 independent replicates per genotype for each reported acylcarnitine species. For box plots in A.-E. and H., center lines show the medians; box limits indicate the 25th and 75th percentiles as determined by R software; whiskers extend 1.5 times the interquartile range from the 25th and 75th percentiles, and outliers are represented by dots. Significance calculated by a two-tailed Student’s t-test; * = p<0.05, ** = p<0.01, *** = p<0.001, n.s. = not significant.

Hypoketotic hypoglycemia is typically associated with defects in fatty acid oxidation (FAO)^29,30^, but can also occur with more generalized mitochondrial defects, such as mtDNA loss^31^. To further explore the underlying cause of this phenotype, we examined metabolites and acylcarnitines in tissues isolated from newborn WT and KO pups. Consistent with the low blood sugar and lactic acidosis seen in knockout serum, metabolomic analysis of liver tissue revealed decreased glucose levels concomitant with increased pyruvate and lactate levels (Figure 2F), suggesting a defect in gluconeogenesis (Figures 2B, D). This analysis also revealed a substantial increase in amino acids whose catabolism occurs within mitochondria, such as branched chain amino acids (BCAAs), in both liver (Figure 2G) and heart (Figure S2F). These data were corroborated via acylcarnitine analysis, as byproducts of BCAA catabolism were all significantly elevated in *Pptc7* knockout liver (Figure 2H, S2H) and heart (Figure S2G, S2I). These acylcarnitine data also suggest widespread defects in FAO, as medium and long chain acylcarnitines were significantly increased in *Pptc7* KO liver and heart tissues, and short chain acylcarnitines levels were increased in heart (Figures 2H, S2G, S2I).

To our knowledge, these *Pptc7* KO-associated metabolic abnormalities are not consistent with any single inborn error of metabolism, but rather share molecular characteristic with various disorders. For instance, glutaric aciduria type II (GAII) has substantial overlap with *Pptc7* KO presentation (hypoketotic hypoglycemia, lactic acidosis, and elevation of multiple acylcarnitines)^32^, yet key distinctions remain (e.g. levels of α-aminoadipic acid, a lysine catabolite, are typically increased in GAII instead of decreased, as is seen in the *Pptc7* KO)^32^. In this way, the pleiotropic effects of Pptc7 disruption on FAO might be most analogous to the established paradigm of “synergistic heterozygosity,” in which combined heterozygous defects in distinct FAO enzymes is sufficient to produce a metabolic phenotype^33^. Overall, these data suggest that Pptc7 function likely influences multiple metabolic pathways, and that its expression is required for the use of multiple nutrients in the perinatal stage.

### Loss of Pptc7 causes a post-transcriptional defect in mitochondrial biogenesis

To profile the molecular consequences of Pptc7 loss, we performed quantitative proteomics on tissues from WT and KO littermates. These results showed no clear pattern of changes in non-mitochondrial proteins for either heart (Figure 3A, S2A) or liver (Figure 3B, S2B), but revealed a widespread decrease of mitochondrial proteins in both tissues (Figures 3C, D, S2A, S2B). These data were corroborated by Western blots of proteins involved in oxidative phosphorylation, which were decreased only in KO heart and liver tissues (Figure S3C, S3D). Notably, Bnip3 – a stress-activated protein involved in mitophagy^34^ – is elevated in KO tissue, suggesting that mitophagy might play a role in these decreased protein levels. Importantly, nuclear-encoded mitochondrial mRNA transcripts were not decreased (other than the CRISPR-targeted *Pptc7*) (Figure 3E), suggesting that disruption of mitochondrial protein homeostasis in *Pptc7*-null tissues occurs post-transcriptionally.

**Figure 3:**
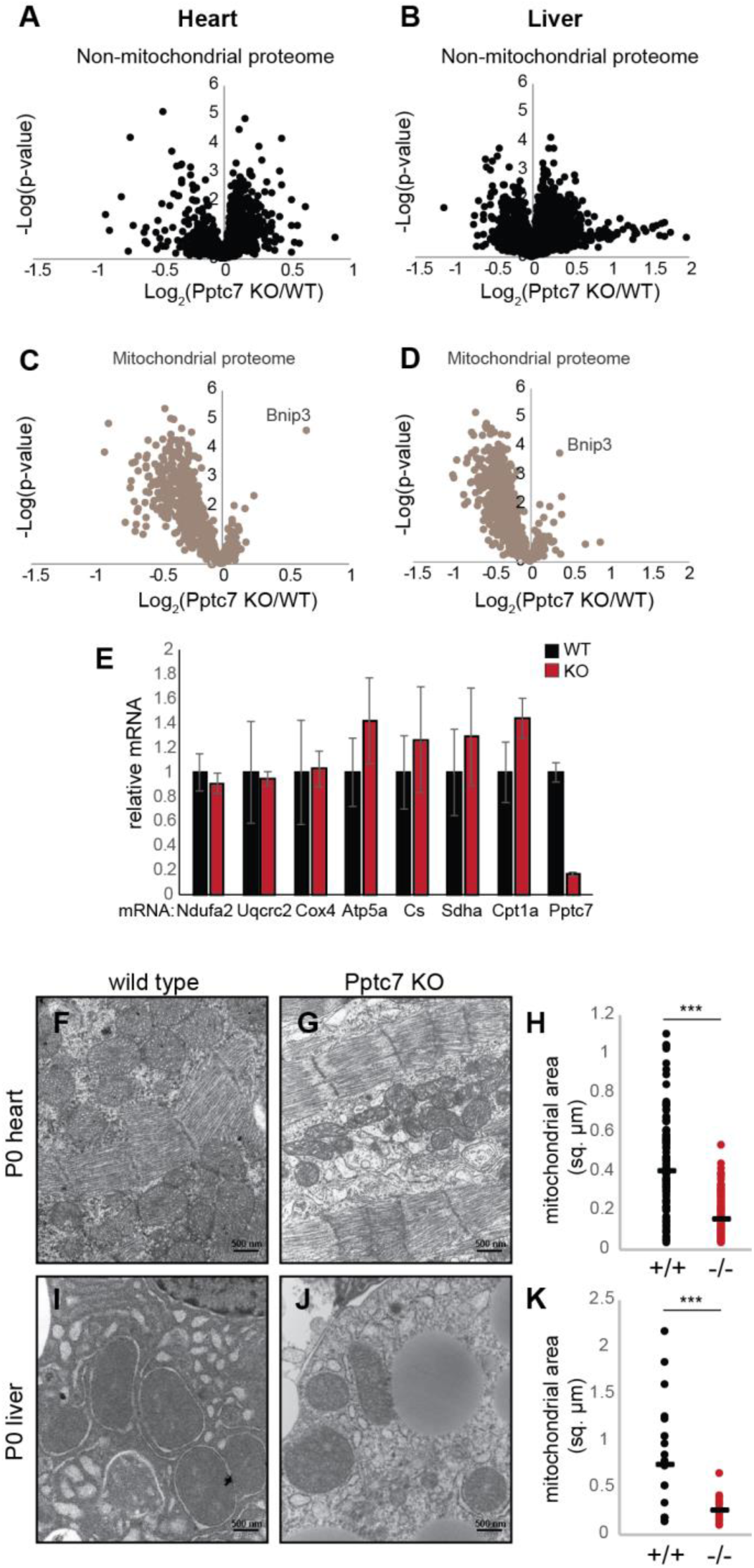
Loss of Pptc7 selectively decreases mitochondrial content. A-D. Volcano plots of non-mitochondrial (A, B) proteins and mitochondrial proteins (C, D) in both heart (A, C) and liver (B, D). One of the only significantly upregulated mitochondrial proteins is Bnip3, a protein upregulated during organellar stress. E. qPCR of select nuclear-encoded mitochondrial targets from WT (black, n=4) or KO (red, n=3) heart tissue shows no significant decrease between genotypes, except for Pptc7. Error bars represent standard deviation; significance calculated with a two-tailed Student’s t-test. F.-H. Transmission electron microscopy (TEM) was used to image heart tissue from WT (F.) and *Pptc7* KO (G.). Mitochondrial area was quantified in ImageJ (one dot = one mitochondrion) using the scale bar (500 nm). KO heart mitochondria are significantly smaller than those in WT tissues (H.). I.-K. TEM images of liver tissue from WT (I.) and Pptc7 KO (J.) and analyzed as described for heart tissue. KO liver mitochondria are significantly smaller than WT mitochondria (K.). For H. and K., each dot represents the area of a single quantified mitochondrion in WT (black) or KO (red) tissues. The line represents the median area in each condition. Significance was calculated using a two-tailed Student’s t-test; *** = p<0.001.

To further examine how this loss of mitochondrial proteins is manifest at the cellular level, we performed electron microscopy of tissue slices from P0 mice. In both heart (Figure 3F, G) and liver (Figure 3I, J) mitochondria from KO mice were significantly smaller than those from WT mice. On average, mitochondria from KO mice had ~40% (Figure 3H) and ~34% (Figure 3K) of the surface area of WT mice in heart and liver, respectively. As such, certain mitochondrial defects that we observe in KO tissues, such as decreased citrate synthase activity (Figure S3E, F) and coenzyme Q levels (Figure S3G, H), likely stem from the overall reduction in mitochondrial content, and not from protein-level regulation; in each case, these changes are no longer apparent when normalized to total mitochondrial protein content (Figure S3E-H). To determine putative Pptc7 substrates whose dysregulation might contribute to these global mitochondrial effects, we next performed phosphoproteomic analysis of WT and KO tissues.

### Phosphoproteomic analysis of *Pptc7*^−/−^ mice reveals candidate substrates

We previously demonstrated that deletion of the yeast ortholog of Pptc7 (termed Ptc7p) caused mitochondrial dysfunction concomitant with elevated phosphorylation on a range of proteins^17^. These results suggest that this phosphatase likely has multiple substrates. We similarly assessed putative substrates for Pptc7 using quantitative multiplexed phosphoproteomic analysis of littermate-matched WT and KO heart and liver tissues (Figure 4A). Analogous to Δ*ptc7* yeast, we find that 28% of mitochondrial phosphorylation events are altered between WT and KO tissues, compared to only 4% of non-mitochondrial phosphoisoforms (Figure S4A). Of mitochondrial phosphoisoforms changing by ≥ 1.5 fold, 98% are elevated (Figure S4B), consistent with the expected outcome of disrupting a mitochondria-specific phosphatase. In heart, 21 mitochondrial phosphoisoforms were elevated by ≥ 1.5-fold with a *p*-value of < 0.05 (Figure 4B, Table 1), seven of which had a Benjamini-Hochberg multiple hypothesis-adjusted q-value < 0.05 (Figure 4C). Similarly, liver exhibited 28 (Figure 4D, Table 1), and eight (Figure 4E) phosphoisoforms meeting these same respective criteria.

**Figure 4:**
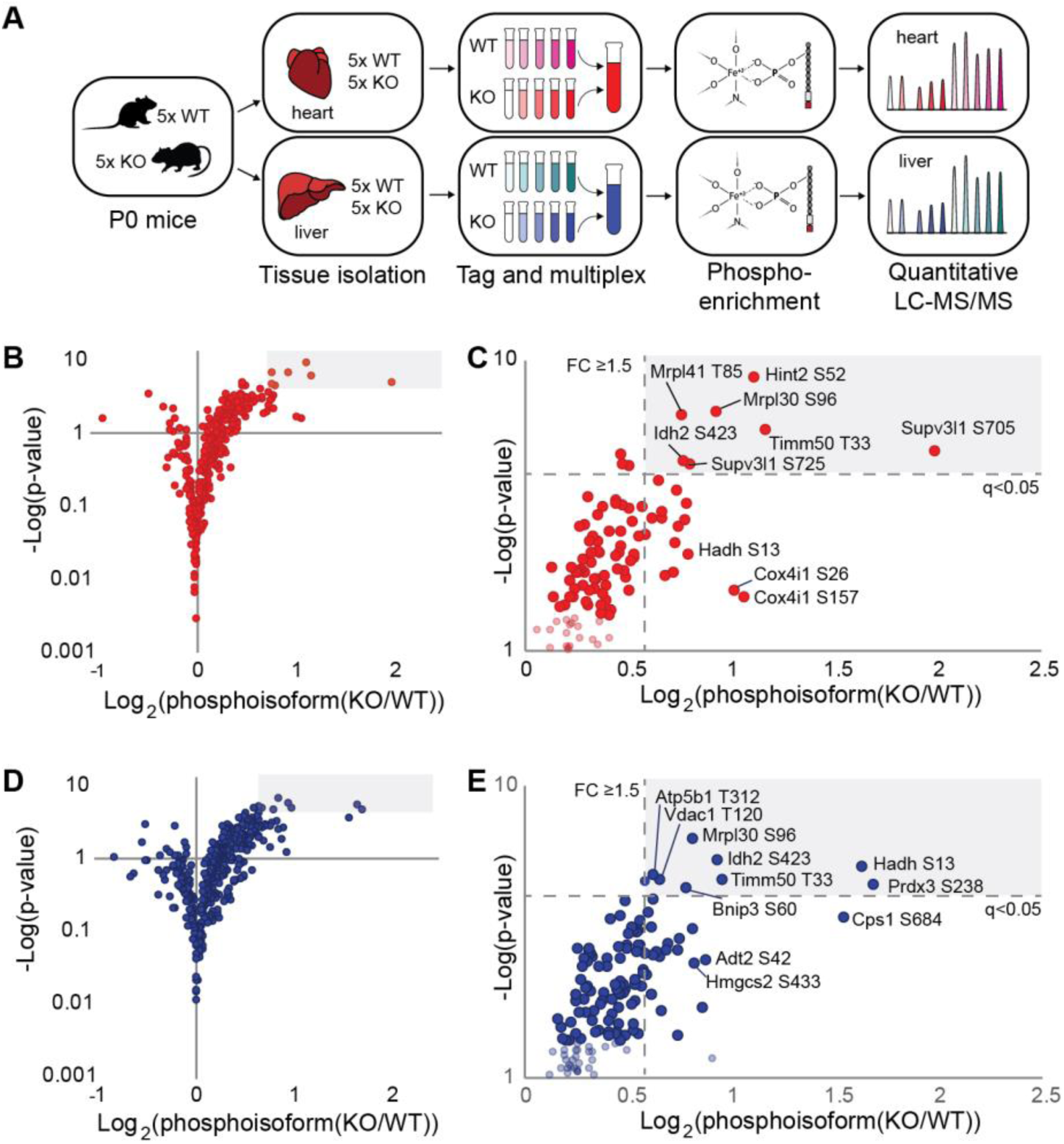
Phosphoproteomic analysis reveals candidate Pptc7 substrates. A. Schematic of multiplexed, quantitative phosphoproteomics for Pptc7 WT and KO heart (red) and liver (blue) tissues. B. Volcano plot showing mitochondrial phosphoisoforms as Log2(fold change) in Pptc7 KO versus WT heart. Shaded region corresponds to identified events with ≥ 1.5 fold change and a q-value of <0.05. D. Zoom in of shaded region in C. showing select statistically significant phosphoisoforms that are candidate Pptc7 substrates. D. Volcano plot showing mitochondrial phosphoisoforms as Log2(fold change) in Pptc7 KO versus WT liver. Shaded region corresponds to identified events with ≥ 1.5 fold change and a q-value of <0.05.E. Zoom in of shaded region in D. showing select statistically significant phosphoisoforms that are candidate Pptc7 substrates.

**Table 1:**
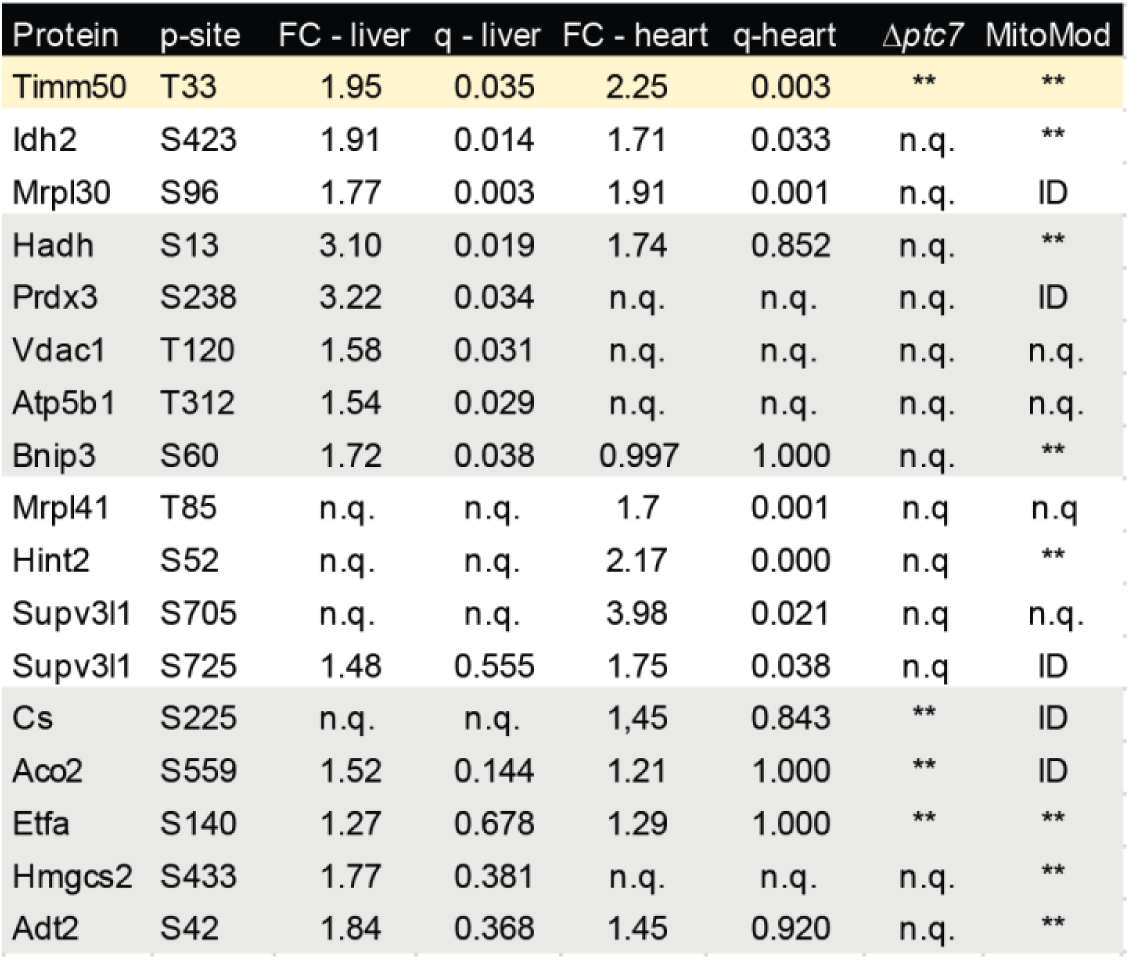
Select Pptc7 candidate substrates across liver, heart, and knockout yeast. Groups of phosphoisoforms shown by protein name and phosphosite identified (p-site), with relative fold changes (FC) and q-values (q) in heart and liver Pptc7 KO tissues. Identification of phosphoproteins in Δ*ptc7* yeast^17^ and in our previous study of obese mice (MitoMod) ^6^ also reported; ** = significantly changing; ID = identified; n.q. = not quantified. Proteins were catalogued into four groups: group 1 (top, white background) are significantly upregulated in both liver and heart KO tissues; group 2 (middle, grey background) are significantly upregulated in liver KO tissue; group 3 (middle, white background) are significantly upregulated in heart tissue; and group 4 (bottom, grey background) notable due to identification in other studies, including in Δ*ptc7* yeast and MitoMod. Timm50 (top, yellow highlight) was significantly altered in both tissues and both previous studies, warranting biological follow up.

Mitochondrial phosphoisoforms elevated in *Pptc7* KO tissues spanned numerous pathways (Table 1), including mitochondrial RNA processing and translation (Mrpl30, Mrps26, Lrpprc, and Supv3l1) (Figure S4C), the TCA cycle (Idh2, Aco2, and Cs) (Figure S4D), and fatty acid oxidation (Hadh and Etfa) (Figure S4E). Notably, we identified overlap between a subset of these elevated phosphoproteins and those found in Δ*ptc7* yeast^17,18^ (Table 1), suggesting that select phosphatase functions are conserved. This broad profile of phosphoproteins is consistent with Pptc7 affecting diverse proteins and pathways, and suggests that it likely influences multiple processes within mitochondria.

### Pptc7-mediated modulation of Timm50 function is conserved through *S. cerevisiae*

We next sought to identify candidate Pptc7 substrates from our analyses above whose dysregulation could give rise to the marked mitochondrial dysregulation we observe in the KO mice. Timm50 is among the most significantly elevated phosphoproteins in *Pptc7* KO tissues, and its identification in both *PTC7* knockout yeast^17^ and our previous study of obese mice^6^ (Table 1) suggests that this phosphorylation may have functional relevance under multiple conditions. The phosphorylation site on Timm50 (Tim50p in yeast) occurs on the matrix-facing N-terminal tail in both the yeast and mouse orthologs (Figure 5A). Although *Timm50* is essential in both *S. cerevisiae^35^* and *M. musculus^36^*, we hypothesized that a less severe disruption of Tim50p/Timm50-mediated protein import activity could generate many of the pleiotropic effects seen in Δ*ptc7* yeast and *Pptc7* KO mice. Indeed, a recent study identified compound heterozygous mutations in *TIMM50* in a patient with a mitochondrial disorder whose tissues exhibited a similar decrease in respiratory chain components as our *Pptc7* KO mice^37^.

**Figure 5:**
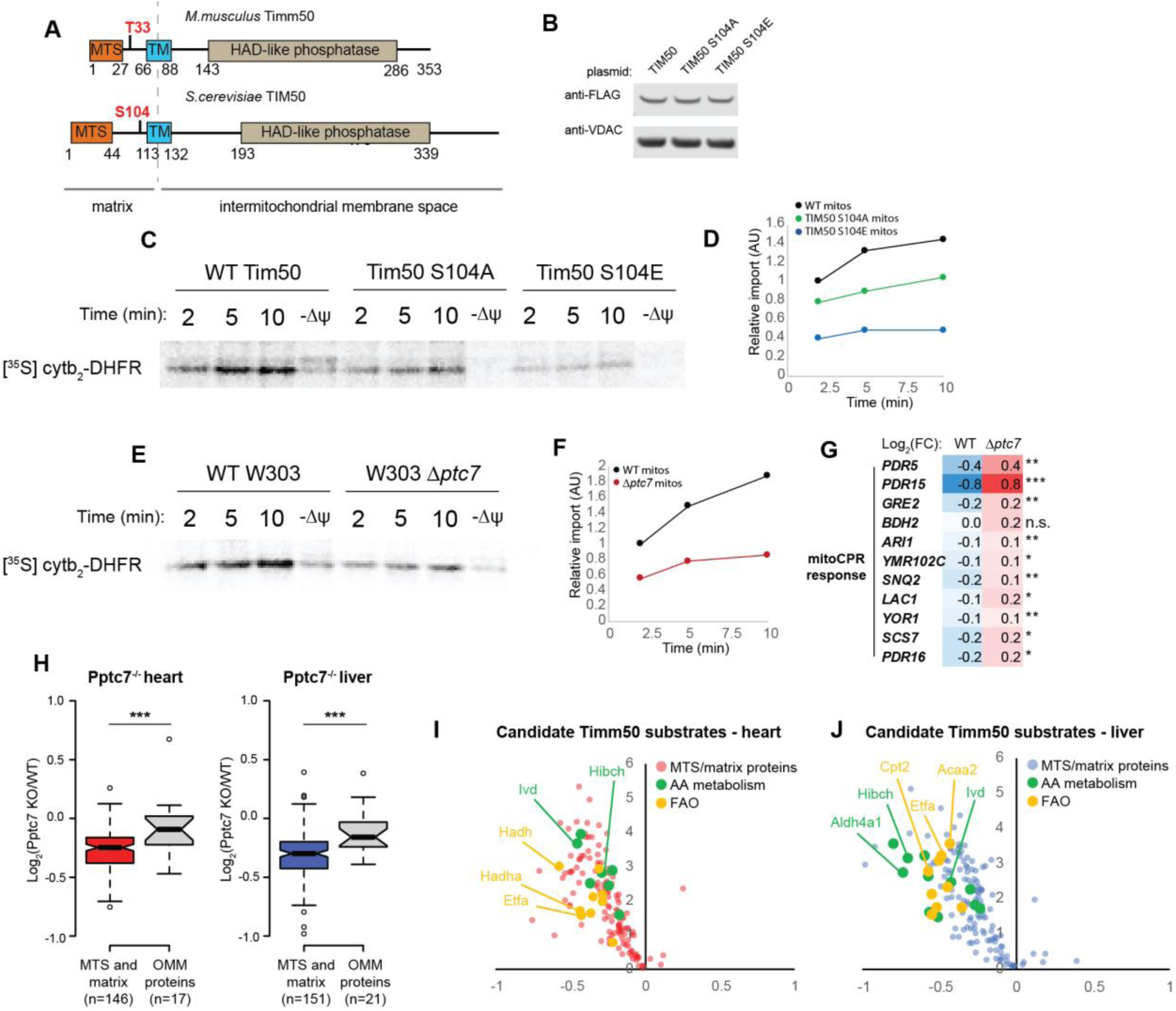
Phosphorylation of the conserved Pptc7 substrate Timm50 decreases mitochondrial protein import in yeast and mice. A. Protein domain schematic of mouse Timm50 (top) and yeast TIM50 (bottom) with highlighted phosphoresidues (red). B. Western blot of overexpressed TIM50 WT, S104A, or S104E shows mutations do not destabilize proteins. C. Import assays in mitochondria isolated from *GAL7*-*TIM50* yeast overexpressing wild type (WT) TIM50 or S104 mutants using the generic import cargo cytochrome b_2_-(167)_ΔG19_-DHFR. D. Quantification of import rates from (C.) E. Import assays in mitochondria isolated from WT or Δ*ptc7* yeast using the matrix-targeted model substrate cytochrome b_2_-(167)_ΔG19_-DHFR. F. Quantification of import rates from (E.). G. Proteomic analysis of WT and Δptc7 yeast show that loss of *ptc7* elevates 11 proteins associated with the mitoCPR response. H. Quantification of mitochondrial proteins in mouse heart (left) and mouse liver (right) showing significantly lower expression protein steady state levels of candidate Timm50 substrates (e.g. MTS and matrix-containing proteins) than select proteins that localize to the outer mitochondrial membrane (OMM). I., J. Volcano plots of candidate Timm50 substrates in heart (I.) and liver (J.) show downregulation of metabolic proteins in amino acid metabolism and fatty acid oxidation, both of which are disrupted in Pptc7-null tissues (Figure 2).

To test whether Tim50p phosphorylation affects mitochondrial protein import, we generated a strain of yeast expressing WT Tim50p, or a non-phosphorylatable (S104A) or phosphomimetic (S104E) FLAG-tagged mutant (Figure 5B). As *TIM50* is essential, we used a previously characterized strain expressing the endogenous *TIM50* gene under a galactose-inducible promoter *(GALT)*^38^, which reduces growth on non-fermentable carbon sources in the absence of galactose (Figure S5A). This phenotype could be rescued fully by exogenous expression of Tim50p (Figure S5A) and Tim50p S104A (Figure S5A), while expression of the phosphomimetic mutant Tim50p S104E exhibited a slight growth defect (Figure S5B). To test whether Tim50p phosphorylation affects import efficiency, we performed import assays using recombinant cytochrome b_2_-(167)_Δ19_-DHFR—a well-established, matrix-targeted model substrate^39^—into mitochondria isolated from strains expressing WT Tim50p or a non-phosphorylatable S104A or phosphomimetic S104E mutant. Mitochondria expressing Tim50p with mutant S104 had import defects, with Tim50p S104E mitochondria importing cytochrome b_2_-(167)_ΔI9_-DHFR almost three-fold less efficiently than WT (Figure 5C, quantified in D). These data suggest that phosphorylation at this site can inhibit mitochondrial protein import.

Given the elevation of Tim50p S104 phosphorylation in Δ*ptc7* yeast^17^, we next tested mitochondrial protein import in Δ*ptc7* yeast and likewise found an approximate three-fold decrease relative to WT import rates (Figure 5E, quantified in F). Δ*ptc7* yeast do not have decreased levels of Tim50p, nor of any other import machinery component^17^ (Figure S5C), suggesting that Tim50p phosphorylation likely contributes to the decreased import rates. Further, proteomic analysis of WT and Δ*ptc7* yeast revealed that the Δ*ptc7* strain exhibits a coordinated upregulation of proteins involved in the mitoCPR—an acute stress response specific to disrupted mitochondrial protein import (Figure 5G)^40^. Collectively, these data suggest that the loss of Ptc7p may elevate Tim50p phosphorylation, dampening mitochondrial import and contributing to a mitoCPR in yeast.

Decreased mitochondrial import may help explain the broad decreases in mitochondrial proteins seen in *Pptc7* KO tissues (Figure 3). Timm50 is a core component of the Timm23 complex that translocates proteins possessing a mitochondrial targeting sequence (MTS) that are typically bound for the matrix^41^. We hypothesized that if Timm50 was selectively impaired in *Pptc7* KO tissues, then matrix-localized MTS-containing proteins would show greater decreases in our proteomics dataset relative to other mitochondrial proteins (e.g. proteins destined for the outer mitochondrial membrane (OMM)). Indeed, proteins that have both an MTS^42^ and matrix localization^43^ were significantly decreased relative to the OMM-localized proteins we measured in both heart and liver (Figure 5H) KO tissues. These decreased proteins include those whose dysfunction is associated with various dysregulated metabolites we identified in *Pptc7* KO tissues: Aldh4a1 (hyperprolinemia)^44^, Pcca, Pccb, Acad8, and Ivd (BCAA and catabolite acylcarnitine accumulation)^45^, and Etfa and Etfdh (medium and long chain acylcarnitine accumulation)^29^ (Figures 5I, J). Together, these data suggest that disruption of Timm50-mediated import could broadly influence metabolic pathways, and may contribute to the widespread defects seen in the *Pptc7* knockout mouse.

### MTS-proximal phosphorylation can influence import and processing

Close examination of our multiple phosphoproteomics datasets revealed a second trend related to mitochondrial protein import. We noticed that amongst elevated (p<0.05) mitochondrial phosphoisoforms identified in mouse and yeast, 11 candidate Pptc7/Ptc7p substrates have phosphorylated residues that lie within or directly proximal to their mitochondrial targeting sequences (Figure 6A). The localization of these phosphorylation events, coupled with data suggesting that Ptc7p/Pptc7 modulates mitochondrial protein import (Figure 5), suggest that the phosphatase may play an addition role in import through the dephosphorylation of these “phospho-MTS” substrates.

**Figure 6:**
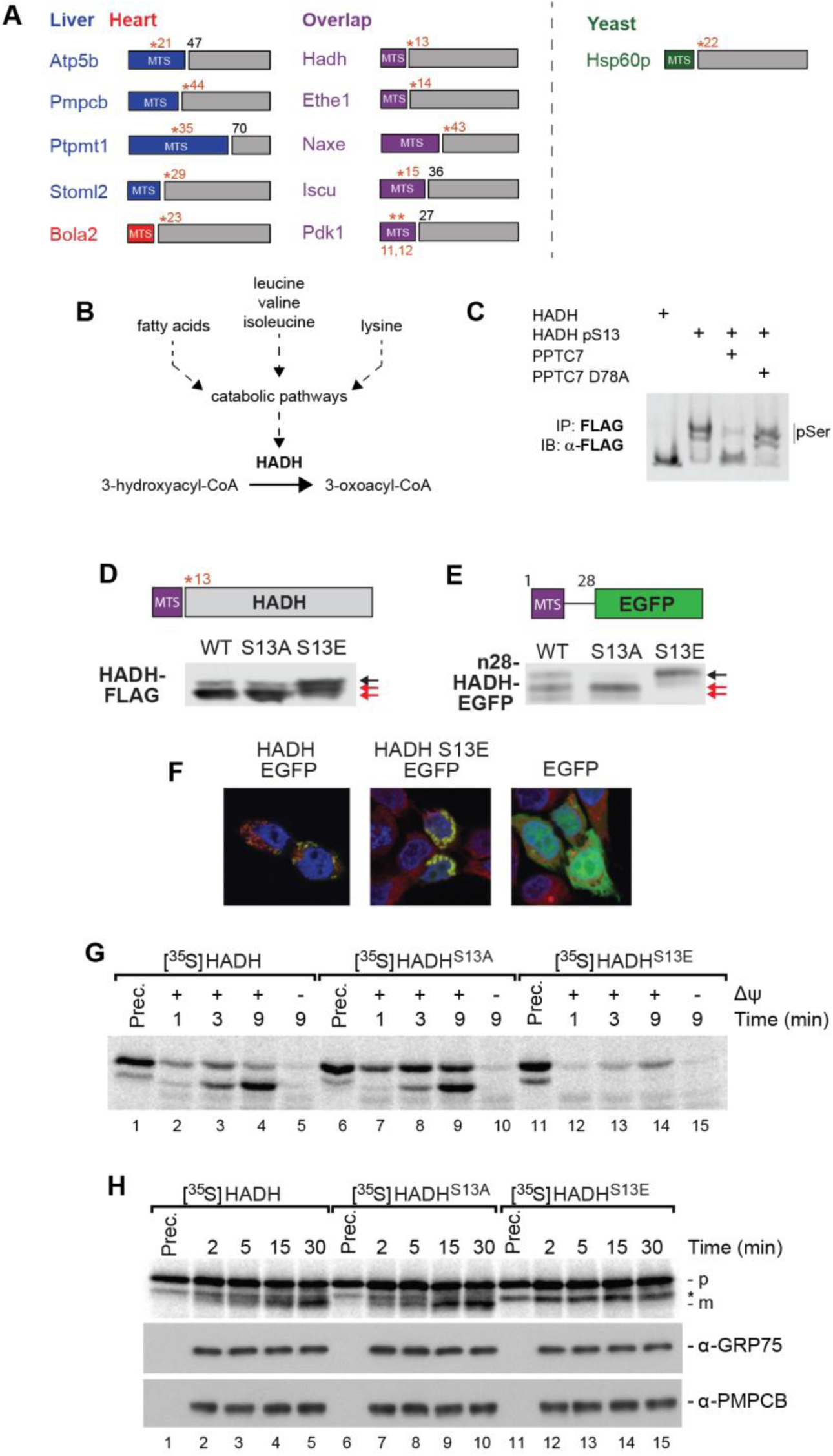
Pptc7 influences mitochondrial protein import and processing through the dephosphorylation of MTS residues. A. Candidate Pptc7 substrates phosphorylated within or proximal to their mitochondrial targeting sequence (MTS) identified in mouse heart (red), mouse liver (blue), and Ptc7p-null yeast (green). B. Hadh is downstream of multiple metabolites dysregulated in Pptc7 knockout mice. C. Recombinant site-specific incorporated HADH pS13 can be dephosphorylated by recombinant PPTC7, but not a catalytically inactive mutant (D78A). D. Overexpression of WT, non-phosphorylatable (S13A) and phosphomimetic (S13E) HADH in 293 cells. S13E causes a shift in HADH processing relative to S13A and WT. E. A fusion protein of the first 28 amino acids of HADH and GFP was made, expressing WT or S13 mutants. S13E is sufficient to disrupt processing of the fusion protein. F. HADH and HADH S13E fusions overlap with MitoTracker Red staining, unlike EGFP with no targeting sequence. G. Import assays with recombinant 35S-labelled HADH, S13A, and S13E show that mutation of S13 alters HADH import efficiency into and processing in mitochondria isolated from HEK293T cells. Prec., precursor protein. H. MPP processing assay shows precursor HADH (p, top band) as well as cleavage into the mature form (m, bottom band) for both wild type and S13A HADH. Processing is completely disrupted in the S13E mutant, shown via the absence of the lowest band.* is a non-specific band in all three conditions. GRP75 and PMPCB blots demonstrate equal loading.

One such phospho-MTS substrate, Hadh, can metabolize multiple nutrients that are disrupted in *Pptc7* KO mice (e.g. BCAAs, lysine, fatty acids)^46^ (Figure 6B), and thus its dysfunction could contributes to the KO metabolic phenotypes. Hadh phosphorylation is also elevated in both tissues, and ranks third in fold change and q-value amongst > 9000 phosphoisoforms identified in liver (Figure 4E, Table 1). To test whether PPTC7 can directly dephosphorylate phospho-MTS substrates, we generated recombinant human HADH with site-specific phosphoserine incorporation at S13^47,48^ (see Methods for details). We validated phosphoserine incorporation using PhosTag gels, which showed a mobility shift for phosphorylated, but not WT, HADH (Figure 6C, lanes 1 and 2). Further, WT PPTC7, but not an inactive mutant, was able to dephosphorylate HADH (Figure 6C). These results were recapitulated with a second phospho-MTS substrate, ETHE1 (Figure S6A), demonstrating PPTC7 can directly dephosphorylate phospho-MTS substrates in vitro.

We next tested whether HADH S13 phosphorylation affects enzyme activity, but found no differences between pS13 and WT HADH (Figure S6B). We reasoned that the proximity of pS13 to the mitochondrial targeting sequence of Hadh may instead affect protein import or processing. Consistently, our proteomics analysis revealed an increased abundance of peptides from precursor Hadh (peptides that span the MTS cleavage site), but not in the mature (i.e., processed) protein (Figure S6C). To further examine Hadh import or post-import protein stability in cells, we expressed phosphomimetic (S13E) and non-phosphorylatable (S13A) HADH in 293 cells and found that the S13E mutant was not properly processed (Figure 6D). We observed similar protein processing and/or stability errors with phosphomimetic mutation of other phospho-MTS substrates, including ETHE1 (Figure S6D), Naxe (Figure S6E), and Iscu (Figure S6F). To determine whether the MTS-proximal phosphomimetic mutation is sufficient to disrupt processing, we generated chimeras containing the N-terminus of HADH fused to GFP and found that the S13E mutation likewise disrupted processing of the fusion protein (Figure 6E, S6G). Importantly, S13E-HADH-GFP still localized to mitochondria (Figures 6F, S6H), suggesting that the processing defect is not due to altered cellular targeting.

We next aimed to measure in vitro mitochondrial import efficiency on phospho-MTS substrates. Our in vitro site-specific phospho-incorporation system noted above could not be imported—an established problem for proteins produced in cell-free and *E. coli*-based systems^49-51^. Instead, we performed import assays with recombinant radiolabeled WT, S13A, and S13E HADH. Each construct could be imported into mitochondria, as evidenced by the time-dependent accumulation of proteinase K-protected species (Figure 6G); however, HADH S13E imported more slowly relative to WT and S13A HADH proteins. Additionally, phosphomimetic mutation of S13 HADH disrupted processing, as only full-length protein accumulated in mitochondria (Figure 6G). To confirm the processing defect, we subjected HADH, S13A, and S13E mutants to an in vitro MPP processing assay that is uncoupled from protein import^52^. MPP (mitochondrial processing peptidase) removes MTS sequences upon entry into the matrix, and this processing event is critical for mature protein stability^53^. We find that phosphomimetic mutation of S13 completely disrupted HADH processing (Figure 6H, bottom band), whereas WT and S13A mutants were processed normally. With the caveat that we are analyzing phosphomimetic mutants, these data suggest that phosphorylation of HADH proximal to its MTS is sufficient to dampen, but not eliminate, mitochondrial import, and that dephosphorylation of HADH after translocation is required for MPP processing and mature protein stability. As failure to remove mitochondrial targeting sequences is associated with disrupted protein stability, these data suggest that an inability to dephosphorylate and subsequently process these proteins may further contribute to the decreased mitochondrial content and metabolic defects seen in Pptc7 KO mice.

## DISCUSSION

Despite numerous studies indicating that a large proportion of mitochondrial proteins are phosphorylated, how these modifications affect organellar function remains unclear. In the present study, we extended our findings on the yeast mitochondrial phosphatase Ptc7p into a mammalian system through the creation and characterization of a *Pptc7* knockout mouse. Surprisingly, we find that *Pptc7* is an essential gene in mammalian development, with its knockout leading to fully penetrant perinatal lethality.

Perinatal lethality in mouse models is often associated with defects in metabolism, when the newborn pup experiences a robust metabolic transition from placental to extrauterine nutrients^23^. *Pptc7* KO pups present with hypoketotic hypoglycemia—typically associated with FAO disorders^29^—which is the likely cause of their death. Despite this presentation, our molecular characterization using metabolomics, proteomics, and electron microscopy suggests a broader mitochondrial dysfunction, which, in certain circumstances, is also associated with hypoketotic hypoglycemia^31^. For example, knockout of *Cluh*, an RNA-binding protein that selectively influences mitochondrial biogenesis, causes perinatal lethality in mice with metabolic defects (e.g. hypoglycemia and hyperprolinemia) coupled with a stark decrease in the mitochondrial proteome^54^. These data suggest that disruption of genes required for general mitochondrial homeostasis (e.g. *Cluh* and *Pptc7*) can cause symptoms similar to classes of inborn errors of metabolism through broad mitochondrial dysfunction.

The global metabolic and mitochondrial defects seen in *Pptc7* KO mice are likely due to aberrantly elevated phosphorylation of mitochondrial proteins. We focused our attention on two subsets of these phosphoproteins that could influence mitochondrial protein import and thus have widespread effects within this organelle. While both subsets affect mitochondrial protein import, we propose that elevated phosphorylation on Timm50 and Hadh mediate mitochondrial dysfunction through two distinct mechanisms: via direct dampening of the import machinery (Timm50), and by decreasing the import rate and/or processing of select mitochondrial-targeted precursor proteins (e.g. Hadh). The mitochondrial import machinery is regulated by kinases in yeast^55-58^, and phosphorylation of TOM complex proteins is plausible given their accessibility to cytoplasmic kinases. Less is known regarding phosphorylation on proteins within the TIM complex, and how the matrix-localized tail of Timm50—a protein spanning the mitochondrial inner membrane—could be accessed by such regulatory molecules is unknown. While a handful of studies suggest protein kinases (e.g. PKA) lie within the mitochondrial matrix^59-61^, it is not well understood how and when these enzymes translocate into the organelle or under what conditions they are active.

One possible mechanism for matrix-localized protein phosphorylation is that the modification of select mitochondrial proteins occurs outside of the mitochondrion. Our data support a model in which phosphorylation outside of mitochondria influences Hadh import rate and downstream protease processing. In plants, phosphorylation of the targeting sequences of chloroplast-bound proteins can alter their import efficiency by promoting or disrupting molecular associations with the import complex itself, or with chaperones that facilitate import^62,63^. Consistently, phosphorylation of mitochondrial precursors such as GSTA4-4^64^, CYP2B1, and CYP2E1^65^ increases their association with cytosolic chaperones (e.g. Hsp70) to promote import, while phosphorylation of the Tom22p precursor in yeast increases its association with the import complex^55^. Alternatively, precursor phosphorylation on mitochondrial proteins including CNP2^66^ and Tom40p^56^ decrease or inhibit targeting to the organelle, consistent with our data on Hadh (Figure 6). Beyond these examples, there are clues that mitochondrial proteins may be phosphorylated by cytosolic kinases on sequences distinct from their MTS: the outer mitochondrial membrane (OMM) protein MitoNEET is phosphorylated within its N-terminus, which may affect its membrane targeting^67^; ferrochelatase is phosphorylated at T116 by OMM-localized PKA to activate enzymatic activity and alter heme biosynthesis^68^, and PKA has also been shown to influence the import of NDUFS4 through phosphorylation of a residue near its C-terminus^69^. Notably, PKA can be anchored to the OMM through a subset of AKAP proteins^70^, suggesting this kinase may be properly localized to phosphorylate mitochondrial-destined proteins. Collectively, these data suggest that cytosolic phosphorylation of mitochondria-bound proteins can affect their import and function through multiple mechanisms. Further, the possibility that phosphorylation affects organellar targeting across species—including yeast, plants, and mammals— lead us to propose that this may be a widespread, but underappreciated, mode of regulation for organelle-bound protein precursors.

Phosphorylation of various organellar targeting sequences across species has been shown to affect association with cytosolic chaperones to enable import. Evidence exists for a similar chaperone to enable mitochondrial import: a biochemically isolated factor, termed mitochondrial import stimulation factor (MSF), was identified as a pair of 14-3-3 proteins^51,71,72^—proteins that preferentially bind to phosphoserine-containing sequences^73^. At least one phospho-MTS target (Iscu pS14) identified in this work binds robustly to 14-3-3 proteins^73^, and this phosphorylation event has been linked to Iscu protein stability^74^. Many phospho-MTS targets have preferred 14-3-3 sequences^75^, suggesting that these phosphorylation events may influence association with MSF, Hsp70^64^, or other chaperones to impact import.

After import into the organelle, dephosphorylation is presumably required for proper processing and maturation. This model was suggested for phosphorylated chloroplast precursor proteins^63^, but a candidate phosphatase has not yet been identified. Plants have multiple *Pptc7* paralogs^76^, with a subset predicted to be chloroplast-localized, suggesting that Pptc7 paralogs likewise may mediate these functions in plants. It is also notable that Timm50 has a vestigial phosphatase domain^77^, which perhaps once functioned to dephosphorylate these proteins before matrix import. In most organisms, Timm50 does not possess critical catalytic residues within its phosphatase domain^77^; however, in select lower organisms, such as the parasitic *T. Brucei*, Timm50 (called TbTim50) retains phosphatase activity^78^. In this species, ablation of catalytic phosphatase residues alters the stability of mitochondrial proteins such as VDAC^78^, suggesting functional relevance. It is tempting to speculate that Timm50, and ultimately Pptc7, have evolved to coordinate dephosphorylation of proteins targeted to the mitochondrion.

Overall, our data add to a growing narrative that mitochondrial PTMs are widespread and impactful. It remains largely unclear whether the bulk of these PTMs are regulatory in nature, or are instead adventitious and unintentionally disruptive, and thus merely need to be removed to maintain optimal mitochondrial function. It is intriguing that our work on phosphorylation draws parallels to mitochondrial acylation, where the enzymes tasked with removing PTMs are better understood and seem to bear the larger metabolic managerial burden. In this regard, it is further noteworthy that mice lacking the mitochondrial deacetylase Sirt3 exhibit similar phenotypes to the Pptc7 KO mice, albeit to a lesser extent and not until they are stressed^79,80^. However, our data also suggest that some mitochondrial phosphorylation events have important regulatory potential. First, the fact Pptc7 KO mice seemingly develop normally, but fail to thrive during a specific stage, is perhaps consistent with a regulatory switch of sorts. Second, our data lead us to speculate that select mitochondrial proteins are phosphorylated in the cytosol, and then dephosphorylated by phosphatases within the organelle. In this model, proteins would be available to various cytosolic kinases, whose classic signaling functions may then serve to direct proteins to mitochondria or to alter their import rates. Regardless of its regulatory potential, our data demonstrate that Pptc7 is required for surviving the perinatal transition, and show that protein dephosphorylation within the mitochondrial matrix is essential for mammalian development. Further studies will be required to understand the full repertoire of Pptc7 substrates, the potential regulation of this phosphatase, and the physiological consequences of Pptc7 disruption in conditions beyond birth.

## METHODS

### Creation of the Pptc7 knockout mouse model

A Pptc7 knockout strain was generated using CRISPR-Cas9 technology in the C57BL/6J (B6) strain of *Mus musculus* [NCBItax:10090]. Mutational details are provided at MGI; Pptc7^em1Pag^ at MGI:6094244, and Pptc7^em2Pag^ at MGI:6143811. The second and third coding exons of Pptc7 were targeted for genome editing using two target sequences to maximize specificity (all predicted off-target sites had (i) at least 3 mismatches, with at least 1 mismatch in the 12bp seed region or (ii) 2 mismatches in the seed region). In vitro transcription template was generated by overlap-extension PCR with one oligo carrying a 5’ T7 adapter, the target sequence, and a portion of the common gRNA sequence, and the other oligo carrying the antisense common gRNA sequence. In vitro template was column-purified and in vitro transcribed with the MEGAshortscript kit (Thermo-Fisher), and the resultant gRNA was cleaned with the MEGAclear kit (Thermo-Fisher). For injection-grade purification, gRNA was ammonium acetate purified, washed with 70% ethanol, and resuspended in injection buffer. One-cell fertilized C57BL/6J embryos derived from mice obtained from Jackson laboratories were microinjected with a mixture of both gRNAs (25 ng/ul each) and Cas9 protein (PNA Bio, 40 ng/ul), and then implanted into pseudopregnant B6 recipients. Tail DNA was harvested from resultant pups at weaning and used for genotyping.

### Breeding, care, and selection of mice for experimental procedures

All animal work was done in accordance with IACUC approval (protocol/animal welfare assurance #A3368-01). The Pptc7^−/−^ *Mus musculus* strain was generated via CRISPR and is registered with MGI under the names C57BL/6J-Pptc7^em1Pag^ (accession number MGI:6094249) and C57BL/6J-Pptc7^em2Pag^ (accession number MGI:6143812); two strains are registered as two independently segregating Pptc7^−/−^ alleles were generated in our founder mouse – one with a 4 bp deletion in exon 2 (hereby called E2; MGI:6094244), and one with a 1 bp deletion in exon 3 (hereby called E3; MGI:6143811). As the phenotypes of all Pptc7-null genotypes were identical (see Figure S1), all genotypes are collectively annotated as “Pptc7^−/−^ mice” and not by their specific alleles unless noted in the text or figure legends. The C57BL/6J wild type strain (Jackson Laboratories) was used for the generation of the Pptc7-CRISPR strain as well as for outbreeding. Mice were housed in a pathogen-free vivarium on a 12-hr light:dark cycle. Mice were group housed by strain and sex under temperature- and humidity-controlled conditions and received ad libitum access to water and food. Upon weaning, mice were maintained on a standard chow diet (Formulab 5008; 17.0% kcal fat; 56.5% carbohydrate; 26.5% protein) Strains were housed within the same vivarium throughout the duration of the study. Studies were performed with P0 mouse pups, sacrificed within 24 hours of their birth unless otherwise noted. Sex was not determined for experiments as males and females are indistinguishable at P0. Littermates were typically selected according to genotype and randomly assigned to experimental groups.

### Genotyping analysis

Tail tips were isolated from each mouse and used as a source of genomic DNA (gDNA). Tails were resuspended in 600 μl of Genomic Lysis Solution (20 mM Tris, pH 8.0, 150 mM NaCl, 100 mM EDTA, 1% SDS) supplemented with 3 μl concentrated proteinase K (Roche) and incubated at 55°C overnight. After the overnight incubation, samples were cooled to room temperature for 10 min. before the addition of 200 μl Protein Precipitation Solution (Qiagen). Samples were then incubated on ice for 5 min. before vortexing for ~15 s. per sample. Samples were centrifuged (16K × g) for 5 min. to pellet precipitated proteins. The supernatant was removed and DNA was purified via isopropanol precipitation. gDNA was quantified using a Nanodrop (Thermo-Fisher) and ~300 ng of gDNA was added to each genotyping reaction. We generated a genotyping strategy that exploited unique restriction sites that were either created (E3d1) or destroyed (E2d4) within each *Pptc7* indel. Within exon 2, the 4 bp deletion disrupted a naturally-occurring BsrBI site, which enabled amplification of ~600 bp flanking the indel and subsequent restriction digest. Within exon 3, the 1 bp deletion generated a PasI site, which enabled amplification of ~330 bp flanking the indel and digestion with PasI (Thermo-Fisher) for 60 min. at 55°C. Products were resolved on a 1% agarose gel.

### Next generation sequencing

The F0 founder mouse was sacrificed at 18 months of age via CO_2_ asphyxiation and tissues isolated, including brain (cerebrum), small intestine, kidney, skeletal muscle (gastrocnemius), spleen, stomach, liver, and heart, and tissues from two heterozygous F1 offspring (age 15 months, female). Genomic DNA (gDNA) was harvested from each tissue using a Qiagen DNeasy Blood & Tissue Kit (Qiagen) according to the manufacturer’s protocol. Each sample was amplified using genotyping primers for E2 or E3 as a template for NGS-compatible sequencing, and then amplified using i5/i7 compatible NGS primers. Purified amplicons were submitted to the University of Wisconsin-Madison Biotechnology Center, where libraries were prepared with guidance from Illumina’s 16s Metagenomic Sequencing Library Preparation Protocol, Part #15044223 Rev. B (Illumina), with slight modifications. Illumina dual indexes and Sequencing adapters were added. Following PCR, libraries were cleaned using a 0.9x volume of AxyPrep Mag PCR clean-up beads, and were standardized to 2 nM and pooled prior to sequencing. Paired end, 150 bp sequencing was performed using the Illumina MiSeq Sequencer and a MiSeq 300 bp (v2) sequencing cartridge. Images were analyzed using the standard Illumina Pipeline, version 1.8.2. Analysis of NGS data was performed by the University of Wisconsin (UW) biotechnology center.

### Serum metabolite measurements

After decapitation of P0 pups, a small amount of whole blood was used to measure blood glucose using a CVS Health Advanced Blood Glucose Meter (CVS). Blood glucose levels were taken at least twice, and averaged values reported for each mouse. The remainder of the blood was collected, spun at 16.1K × *g* for 10 min. at room temperature, serum isolated and stored at -80°C until use. Circulating insulin was assayed using the Ultra Sensitive Mouse Insulin ELISA kit (Crystal Chem) according to manufactuer’s instructions using the “Low Range Assay” protocol. Briefly, serum from n=8 WT, n=5 heterozygous, and n=10 KO mice was thawed on ice. 5 μl of serum was assayed for each mouse and quantified relative to the standard curve generated from the kit. Lactate levels were measured in serum isolated from n=9 WT, n=20 heterozygous, and n=7 KO mice mice using the Lactate Colorimetric/Fluorometric Assay Kit (BioVision) according to manufacturer’s instructions, using 5 μl of serum from each mouse quantified relative to the standard curve generated from the kit. Ketones were measured in serum isolated from newborn mice using the Ketone Bodies Kit (both R1 and R2 sets, Wako Biosciences) as previously described^6^. Serum from n=9 WT, n=13 heterozygous, and n=6 KO mice was thawed on ice, and 5 μl was assayed for each mouse and quantified relative to a β-hydroxybutyrate standard curve. For all serum tests, values reported are averaged from all measurements per group, and significance was tested using a two-tailed Student’s t-test.

### Metabolomics

Heart and liver tissues were immediately harvested and snap frozen in liquid nitrogen unless otherwise noted. Metabolites were extracted from cryo-pulverized tissue with a solvent mixture consisting of 7:2:1 HPLC grade methanol:water:chloroform; this was followed by a subsequent extraction of lipids by addition of chloroform to 50% of final volume. Acylcarnitines were analyzed by reversed phase liquid chromatography – tandem mass spectrometry on a Q Exactive Focus operating in positive ion mode (RPLC-ESI-MS/MS). Amino acids and other polar metabolites were derivatized with methoxyamine-HCl and MSTFA and analyzed by gas chromatography – electron ionization mass spectrometry on a GC-Orbitrap (GC-EI-MS). Lipids and Co-enzyme Q (CoQ) intermediates were analyzed by RPLC-ESI-MS/MS in alternating positive and negative ion mode, with PRM targets for selected CoQ intermediates^81^. Acyl-carnitines and CoQ intermediates were quantified from peak area using Thermo’s Tracefinder application. GC features were quantified using in-house software (modified from ^82^), and lipid features were quantified using the Thermo’s Compound DiscovererTM 2 application. Identification of features was performed with spectra matching to library of standards (NIST 14 ^83^) or predicted fragmentation spectra (LipiDex, ^84^).

### Electron microscopy

Tissues were harvested from litters of live P0 pups and were quickly washed in PBS and fixed in ~5 ml of fixation buffer (2.5% glutaraldehyde, 2.0% paraformaldehyde in 0.1M sodium phosphate buffer, pH 7.4) overnight @ 4°C. The tissue was then post fixed in 1% Osmium Tetroxide in the same buffer for 3 hrs @ RT, and the samples were dehydrated in a graded ethanol series, then further dehydrated in propylene oxide and embedded in Epon epoxy resin. Semi-thin (1 μm) sections were cut with a Leica EM UC6 Ultramicrotome and collected on 200 mesh copper grids. Sections were and contrasted with Reynolds lead citrate and 8% uranyl acetate in 50% EtOH. Ultrathin sections were observed with a Philips CM120 electron microscope and images were captured with an AMT BioSprint side mounted digital camera using AMT Capture Engine software.

### Enzyme assays

Citrate synthase activity was assayed as previously described^17^ using genotype verified tissues (n=3-4 each genotype, wild type and Pptc7 knockout heart and tissues). Error was calculated using standard deviation, and significance was calculated using a Student’s t test. Hadh activity was assayed as previously described^85^. Briefly, recombinant human HADH (Entrez Gene #3033) or HADH site-specifically phosphorylated at S13 (see “Phosphoserine incorporation of recombinant proteins using cell free protein synthesis” for more details) were generated with a C-terminal FLAG tag using cell free protein synthesis. Proteins were immunoprecipitated (IPed) using M2-FLAG antibody-conjugated magnetic beads, eluted in FLAG peptide, and HADH activity was assayed using 2.5 μl eluate from each IP (corresponding to ~200 ng recombinant protein per reaction). Error was calculated using standard deviation, and significance was calculated using a Student’s t test.

### Quantitative multiplexed proteomics and phosphoproteomics

Protein from lysed, homogenized mouse tissues was digested into tryptic peptides following previously reported protocols.^86^ This yielded sufficient material to label 0.35 mg of each heart sample and 0.5 mg of each liver sample with tandem mass tags (TMT10plex Isobaric Label Reagent Set, Thermo-Fisher). Labeling was performed following manufacturer recommended protocols except for the peptide:label ratio, which was changed to 0.5:0.8::mg:mg. Equal amounts of sample were combined for each 10-plex experiment following validation of labelling efficiency (> 95% labelling of N-terminal amines). Phosphopeptide enrichment with titanium chelation (Ti-IMAC, ReSyn Biosciences) was performed on pooled samples using previously reported methods ^87^. The depleted sample was saved for protein quantitation. 500 μg of each depleted sample and the entirety of each phosphopeptide enrichment were separated over an Acquity BEH C18 reverse phase column (130 Å pore size, 1.7 μm particle size, 2.1 × 100 mm, Waters Corp) held at 60 °C using a Dionex Ultimate 3000 uHPLC (600 μL/min flow rate, Thermo-Fisher) running with basic mobile phases. Resulting fractions were analyzed on a q-LTQ-OT hybrid mass spectrometer (Orbitrap Fusion Lumos, Thermo-Fisher) operated under data dependent acquisition following nano-LC separation and electrospray ionization. MS1 survey scans were performed in the orbitrap (60K resolution, AGC – 1e6, 50 ms max injection time). Product ion scans following HCD fragmentation (35% NCE) were also performed in the orbitrap (60K resolution, AGC – 2e5, 118 ms max injection time). Monoisotopic precursor selection and dynamic exclusion (60 s) were enabled. Thermo RAW files were searched with the Open Mass Spectrometry Search Algorithm (OMSSA) within the Coon OMSSA Proteomic Analysis Software Suite (COMPASS) against a database of canonical proteins and isoforms (Uniprot, *Mus musculus*) ^88,89^. TMT labelling was imposed as a fixed modification at lysines and N-termini and variable at tyrosines. Additionally, phosphorylation of serine, threonine, and tyrosine, as well as accompanying neutral losses, were set as variable modifications for enriched samples. Search results were filtered to a false discovery rate of 1% at the peptide and protein levels. Sites of phosphorylation were considered localized if given a localization score >0.75 by the PhosphoRS module within COMPASS ^90^. Phosphopeptide intensities were normalized to the total reporter ion intensity at the protein level, as well as, protein mean-normalized fold change. An associated P value was calculated using Student’s t test assuming equal variance. Multiple hypothesis testing was performed by Benjimini-Hochberg correction.

### RNA isolation and qPCR

RNA was isolated using an RNeasy kit (Qiagen) per manufacturer’s instructions. cDNA was prepared using 500 ng of total RNA using the SuperScript III First-Strand Synthesis System (Thermo-Fisher). cDNA was made exclusively with the oligo(dT) primer, and the kit was used according to the manufacturer’s protocol. cDNA was normalized, and 100 ng was used as a template for each qPCR reaction. Reactions were set up using 1× PowerSYBR master mix (Thermo-Fisher), 200 nM forward and reverse primers (each) and water to a final volume of 20 μl total. qPCR was run on a QuantStudio 6 Flex (Applied Biosystems) and data were captured using QuantStudio Real Time PCR system software. Data were analyzed using the ΔΔCt method (n=3-4 samples per genotype), with error bars representing standard deviation and significance calculated using a two-tailed Student’s t test.

### Cloning and site directed mutagenesis

*Escherichia coli* strain DH5α (NEB) was used for all cloning applications and grown at 37 °C in LB media with antibiotics. *Escherichia coli* strain BL21-CodonPlus (DE3)-RIPL (Agilent) was used for all protein expression and purification purposes. Site directed mutagenesis was performed as previously described ^81^.

### Mitochondrial import assays

Various strains derived from the haploid W303 (*his3 leu2 lys2 met15 trp1 ura3 ade2*) yeast were used for mitochondrial isolation for import assays. The Δ*ptc7* strain was generated in W303 yeast as previously described ^17^. The Tim50p studies were performed with a previously described W303 strain in which the *TIM50* ORF was placed under control of the *GAL7* promoter ^38^. For mitochondrial isolation from yeast, a single colony of W303 yeast was incubated in 3-4 ml of YPD media overnight. 1 × 10^8^ cells were diluted into 500 ml of YEP supplemented with 3% glycerol 0.1% dextrose. Yeast were grown at 30°C in an orbital shaker (230 rpm) for 19-20 hrs to a final OD of ~5-6 (with an OD of 1 = 1e7 cells). Mitochondria were isolated as previously described for mitochondrial import ^91^. Isolated mitochondria were resuspended using wide-bore tips to a final concentration of 10 mg/ml in SEM buffer (250 mM sucrose, 1 mM EDTA, and 10 mM MOPS, pH 7.2), snap frozen in liquid nitrogen, and stored at -80°C until use in import assays. Recombinant HADH or cytochrome b_2_-(167)_Δ19_-DHFR (a gift from Elizabeth Craig) were generated using the quick TnT^®^ Quick Coupled Transcription/Translation System (Promega) using either the T7 kit (HADH) or the SP6 kit (cytochrome b_2_-(167)_Δ19_-DHFR) supplemented with ^35^S-Methionine and Cysteine (EasyTag EXPRESS35S protein labeling mix, Perkin Elmer) according to the manufacturer’s instructions. Reactions were terminated by placing reactions on ice until use in mitochondrial import assays. For mammalian targets, radiolabeled HADH (WT, S13A, S13E) precursor proteins were generated by in vitro transcription/translation reactions in rabbit reticulocyte lysate in the presence of ^35^S-Methionine. Mitochondria were isolated from HEK293T cells as described before ^92^. For import assays, mitochondria were resuspended in import buffer (250 mM sucrose, 5 mM magnesium acetate, 80 mM potassium acetate, 10 mM sodium succinate, 20 mM HEPES-KOH, pH 7.4) supplemented with 1 mM DTT and 5 mM ATP. Where indicated, the membrane potential was dissipated prior to the import reaction by addition of 8 μM antimycin A, 1 μM valinomycin and 20 μM oligomycin (AVO). Import reactions were started by addition of radiolabeled precursor proteins followed by incubation at 37°C for indicated times. Reactions were stopped by addition of AVO and non-imported precursor proteins removed by digestion with Proteinase K. Mitochondria were re-isolated by centrifugation at 10,000 × g for 10 min at 4°C and washed once in import buffer. Samples were analyzed on SDS-PAGE followed by digital autoradiography. Images were quantified using ImageJ.

### MPP processing assay

MPP processing assays using soluble mitochondrial extracts from mouse liver were performed as described previously ^52^. In short, mitochondria were lysed in digitonin and the obtained extract incubated with radiolabeled precursor proteins. Samples were incubated for various time points at 37°C and reactions stopped by addition of 4x Laemmli buffer containing 2% (v/v) β-mercaptoethanol. Samples were analyzed by SDS-PAGE and blotted onto PVDF membranes followed by autoradiography and immunodetection.

### Generation of cell free protein synthesis reagents for phosphoserine incorporation

Cell free protein synthesis (CFPS) was used to generate phosphoserine incorporation as previously described ^47^. The bacterial strain C321.ΔA.Aserb.Δmp^S^ was a gift from Jesse Rinehart (Addgene plasmid #68306) and was used to generate lysates for the CFPS reactions as described ^47^ Bacteria were grown in 2xYPTG media (16g/L tryptone, 10g/L yeast extract, 5g/L NaCl, 5g/L K_2_HPO_4_, 3g/L KH_2_PO_4_, and 18g/L glucose filter-sterilized) with 2 mM phosphoserine and antibiotic selection at 30 °C, 220 rpm. At OD 0.6, 1 mM IPTG was added and, at OD 3, cells were harvested by centrifugation (5,000 rcf, 15 min, 4 °C). Bacteria were washed 3 times in cold S30 buffer (10 mM Tris-acetate pH 8.2, 14 mM magnesium acetate, 60 mM potassium acetate) with 1 mM DTT and the pellet was weighed, snap frozen, and stored at -80°C until lysis. Thawed bacteria were lysed by resuspension in S30 buffer with 2 mM DTT (0.8 mL per 1 g wet cells) and sonicated as described by Kwon and Jewett ^93^ (~900 J per 1 L cells). Lysates were supplemented with 1 mM DTT, clarified by centrifugation (12,000×g, 10 min, 4 °C), and the clarified supernatant was incubated in a run-off reaction (1 hour, 250 rpm, 37 °C, covered in foil). The run-off reaction was clarified by a second centrifugation (12,000×g, 30 min, 4 °C) and supernatant was collected with careful avoidance of loose cell debris. Lysates were aliquoted, snap frozen, and stored at -80°C until use.

### Phosphoserine incorporation of recombinant proteins using cell free protein synthesis

CFPS reactions were performed as described by Oza et al^47^. Reactions contained 30% v/v cell extract supplemented with 1.2 mM ATP*, 0.85 mM each of GTP*, UTP*, and CTP* (pH 7 – 7.2); 34 μg/mL folic acid*; 170 μg/mL of *E. coli* tRNA mixture*; 13.3 μg/mL plasmid (maxiprepped); 100 μg/mL T7 RNA polymerase; 2 mM each of standard amino acids* (omitting methionine and cysteine in S35 experiments); 0.33 mM NAD*; 0.27 mM coenzyme A*; 1.5 mM spermidine*; 1 mM putrescine*; 4 mM oxalic acid*; 130 mM potassium glutamate^‡^; 10 mM ammonium glutamate‡ 12 mM magnesium glutamate^‡^; 33 mM phosphoenolpyruvate (pH 7), 2 mM phosphoserine, 57 mM HEPES pH 7*.

Some reagents can be pre-mixed, aliquoted, and stored at -20 °C (*pre-mix, †amino acid mix prepared as described ^94^; ^‡^salt mix at pH 7 with KOH) and the final reaction should be at pH 6.5-7. CFPS reactions were initiated by thoroughly mixing the cell extract, incubated for 17-20 hours at 30 °C, and performed in 15, 30, or 50 uL reactions. For 50 uL S35 experiments, reactions were performed in a parafilmed, screwcap microcentrifuge tube with shaking (350 rpm).

### PPTC7 and Ptc7p recombinant protein purification and phosphatase assays

N41-Ptc7p and catalytically inactive mutants were prepared as previously described ^17^. HiS8-MBP-tev-PPTC7^NΔ32^ and mutants were expressed in *E. coli* (BL21[DE3]-RIPL strain) by autoinduction as previously described ^17^. Cells were isolated and resuspended in lysis buffer (100 mM HEPES pH 7.2, 300 mM NaCl, 5 mM BME, 0.25 mM PMSF, 1 mg/mL lysozyme (Sigma), and 7.5% glycerol) Cells were lysed by sonication (4 °C), clarified by centrifugation (15,000 *g*, 30 min, 4 °C), and mixed with cobalt IMAC resin (Talon resin) for one hour (4 °C). Resin was washed with Wash Buffer (20 bed volumes: Lysis buffer without lysozyme) and His-tagged protein was eluted with Elution Buffer (50 mM HEPES pH 7.2, 150 mM NaCl, 5 mM BME, 5% glycerol 100 mM imidazole). Eluted protein was concentrated with a 50-kDa MW-cutoff spin filter (Merck Millipore Ltd.) and exchanged into Storage Buffer (50 mM HEPES pH 7.2, 150 mM NaCl, 5 mM BME, 5% glycerol). His8-MBP-PPTC7^NΔ32^ was incubated with TEV protease (1:50, TEV/fusion protein, mass:mass using ε_280_ = 88,615 M^−1^cm^−1^ and MW = 73.9 kDa) for 1 hour (25 °C), then incubated with cobalt IMAC resin for 1 hour (4 °C). Cleaved PPTC7^NΔ32^ (ε_280_ = 20,775 M^-1^cm^-1^, MW = 29.6 kDa) was collected, concentrated with a 10-kDa MW-cutoff spin filter, and exchanged into Storage Buffer. Protein was aliquoted, frozen in N2, and stored at -80 °C. Phosphatase assays were run in 10 μl total volume comprised of 50 mM Tris, pH 8.0, 5 mM MnCl2, 5 μl of site-specifically phosphorylated recombinant protein (~400 ng substrate) and ~200 ng active (wild type) or catalytically inactive (D/A mutants) phosphatase per reaction. Reactions were allowed to proceed for 30 minutes at room temperature, after which they were terminated through the addition of sample buffer to 1×, boiled at 95°C, and run on PhosTag gels to determine dephosphorylation efficiency.

### Transfection and protein expression in 293 cells

293 cells were cultured in DMEM (high glucose, no pyruvate, Thermo-Fisher) supplemented with 10% heat inactivated fetal bovine serum (FBS) and 1x penicillin/streptomycin. Cells were grown in a temperature-controlled CO2 incubator at 37°C and 5% CO2. Cells were subcultured using 0.05% trypsin-EDTA every 2-3 days. For transfections, cells were split to ~40% confluence on Day 1. On Day 2, cells were transfected with 7.5 μg Maxiprep-purified plasmid supplemented with 20 μlg linear polyethylenimine (PEI, PolySciences), and 900 μL Opti-MEM (LifeTechnologies) as previously described ^95^ After 48 hours, cells were collected or fixed for downstream applications.

### Fluorescence Microscopy

Glass cover slips (#1.5) were sterilized and placed into single wells of 12 well plates. 293 cells were seeded at ~60% confluence and transfected with plasmids encoding GFP or N-terminal HADH-GFP fusions as previously described ^95^ 20 hours after transfection, cells were labeled with 25 nM MitoTracker Red CMXRos for 15 minutes at 37°C. Cells were then washed 3× with PBS, and fixed using 4% paraformaldehyde in PBS for 15 minutes at room temperature. After fixation, cells were washed 3× with PBS, and nuclei were stained for 5 minutes with 2 μg/ml [final c] of Hoechst. After nuclear staining, cells were washed again 3× with PBS and mounted to glass slides using ProLong Diamond reagent overnight in the dark, per manufacturer’s instructions. Cells were imaged on a Nikon A1R-SI+ confocal microscope using a 60× oil-based objective using constant settings (e.g. laser intensity) across all images acquired. Images were collected in all three fluorescent channels using NIS element software.

### Quantification and Statistical Analysis

See each individual method for the associated statistical analysis.

## ACKNOWLEDGEMENTS

We thank members of the Pagliarini laboratory for helpful discussions and Amy Lin for her assistance with figure generation. The authors thank the Genome Editing and Animal Modeling core at the University of Wisconsin Biotechnology Center, particularly Kathy Krentz and Dustin Rubinstein, for their design and creation of the CRISPR-Cas9 Pptc7 knockout model. We also thank the University of Wisconsin Biotechnology Center DNA Sequencing Facility for providing next generation sequencing (NGS) services, and the UWBC Bioinformatics Resource Center for the analysis of the NGS data. We thank Ben August and the UW Electron Microscope (EM) Facility for processing samples and for providing training and expertise in EM data acquisition and analysis. We thank Benjamin Des Soye and Michael Jewett for protocols and assistance with the cell free protein synthesis (CFPS) experiments. We thank Elizabeth Craig and her laboratory for the cytochrome b_2_-(167)_Δ19_-DHFR plasmid, advice, and protocols on mitochondrial import assays. We thank Toshiya Endo for the kind gift of the *GAL7::TIM50* strain used in the study. Research reported in this publication was supported by the National Institute of Diabetes and Digestive and Kidney Diseases and National Institute of General Medical Sciences of the National Institutes of Health under award numbers R01DK098672 (to D.J.P.), T32DK007665 (to N.M.N.), and P41GM108538 (to J.J.C. and D.J.P). The Genome Editing and Animal Modeling core at UW is supported by a University of Wisconsin Carbone Cancer Center Support Grant (P30 CA014520). This work was further supported by a Morgridge Postdoctoral Research Fellowship (to K.A.O.), the Deutsche Forschungsgemeinschaft and the Excellence Initiative of the German Federal & State Governments (EXC 294 BIOSS) (to C.M.), and the Emmy-Noether Programm of the Deutsche Forschungsgemeinschaft (to F.N.V).

## AUTHOR CONTRIBUTIONS

N.M.N. and D.J.P conceived of the project and its design and wrote the manuscript. N.M.N. and K.L.S. maintained the mouse colony. N.M.N., F.N.V., D.C.L. prepared samples and performed biochemical experiments. G.M.W. and K.A.O. acquired mass spectrometry data. N.M.N., G.M.W., K.A.O., F.N.V., D.C.L., A.D.A., C.M., J.J.C. and D.J.P. analyzed data.

## REFERENCES

1 Nunnari, J. & Suomalainen, A. Mitochondria: in sickness and in health. Cell 148, 1145-1159, doi:10.1016/j.cell.2012.02.035 (2012).

2 Covian, R. & Balaban, R. S. Cardiac mitochondrial matrix and respiratory complex protein phosphorylation. Am J Physiol Heart Circ Physiol 303, H940-966, doi:10.1152/ajpheart.00077.2012 (2012).

3 Pagliarini, D. J. & Dixon, J. E. Mitochondrial modulation: reversible phosphorylation takes center stage? Trends Biochem Sci 31, 26-34, doi:10.1016/j.tibs.2005.11.005 (2006).

4 Trub, A. G. & Hirschey, M. D. Reactive Acyl-CoA Species Modify Proteins and Induce Carbon Stress. Trends Biochem Sci 43, 369-379, doi:10.1016/j.tibs.2018.02.002 (2018).

5 Carrico, C., Meyer, J. G., He, W., Gibson, B. W. & Verdin, E. The Mitochondrial Acylome Emerges: Proteomics, Regulation by Sirtuins, and Metabolic and Disease Implications. Cell Metab 27, 497-512, doi:10.1016/j.cmet.2018.01.016 (2018).

6 Grimsrud, P. A. et al. A quantitative map of the liver mitochondrial phosphoproteome reveals posttranslational control of ketogenesis. Cell Metab 16, 672-683, doi:10.1016/j.cmet.2012.10.004 (2012).

7 Hebert, A. S. et al. Calorie restriction and SIRT3 trigger global reprogramming of the mitochondrial protein acetylome. Mol Cell 49, 186-199, doi:10.1016/j.molcel.2012.10.024 (2013).

8 Yadava, N., Potluri, P. & Scheffler, I. E. Investigations of the potential effects of phosphorylation of the MWFE and ESSS subunits on complex I activity and assembly. Int J Biochem Cell Biol 40, 447-460, doi:10.1016/j.biocel.2007.08.015 (2008).

9 Linn, T. C., Pettit, F. H. & Reed, L. J. Alpha-keto acid dehydrogenase complexes. X. Regulation of the activity of the pyruvate dehydrogenase complex from beef kidney mitochondria by phosphorylation and dephosphorylation. Proc Natl Acad Sci U S A 62, 234-241 (1969).

10 Dittenhafer-Reed, K. E. et al. SIRT3 mediates multi-tissue coupling for metabolic fuel switching. Cell Metab 21, 637-646, doi:10.1016/j.cmet.2015.03.007 (2015).

11 Wagner, G. R. & Hirschey, M. D. Nonenzymatic protein acylation as a carbon stress regulated by sirtuin deacylases. Mol Cell 54, 5-16, doi:10.1016/j.molcel.2014.03.027 (2014).

12 Phillips, D., Aponte, A. M., Covian, R. & Balaban, R. S. Intrinsic protein kinase activity in mitochondrial oxidative phosphorylation complexes. Biochemistry 50, 2515-2529, doi:10.1021/bi101434x (2011).

13 Weinert, B. T., Moustafa, T., Iesmantavicius, V., Zechner, R. & Choudhary, C. Analysis of acetylation stoichiometry suggests that SIRT3 repairs nonenzymatic acetylation lesions. EMBO J 34, 2620-2632, doi:10.15252/embj.201591271 (2015).

14 Wu, R. et al. A large-scale method to measure absolute protein phosphorylation stoichiometries. Nat Methods 8, 677-683, doi:10.1038/nmeth.1636 (2011).

15 Baeza, J., Smallegan, M. J. & Denu, J. M. Mechanisms and Dynamics of Protein Acetylation in Mitochondria. Trends Biochem Sci 41, 231-244, doi:10.1016/j.tibs.2015.12.006 (2016).

16 Pagliarini, D. J. et al. A mitochondrial protein compendium elucidates complex I disease biology. Cell 134, 112-123, doi:10.1016/j.cell.2008.06.016 (2008).

17 Guo, X. et al. Ptc7p Dephosphorylates Select Mitochondrial Proteins to Enhance Metabolic Function. Cell Rep 18, 307-313, doi:10.1016/j.celrep.2016.12.049 (2017).

18 Guo, X., Niemi, N. M., Coon, J. J. & Pagliarini, D. J. Integrative proteomics and biochemical analyses define Ptc6p as the Saccharomyces cerevisiae pyruvate dehydrogenase phosphatase. J Biol Chem 292, 11751-11759, doi:10.1074/jbc.M117.787341 (2017).

19 Damuni, Z., Merryfield, M. L., Humphreys, J. S. & Reed, L. J. Purification and properties of branched-chain alpha-keto acid dehydrogenase phosphatase from bovine kidney. Proc Natl Acad Sci U S A 81, 4335-4338 (1984).

20 Martin-Montalvo, A. et al. The phosphatase Ptc7 induces coenzyme Q biosynthesis by activating the hydroxylase Coq7 in yeast. J Biol Chem 288, 28126-28137, doi:10.1074/jbc.M113.474494 (2013).

21 Awad, A. M. et al. Chromatin-remodeling SWI/SNF complex regulates coenzyme Q6 synthesis and a metabolic shift to respiration in yeast. J Biol Chem 292, 14851-14866, doi:10.1074/jbc.M117.798397 (2017).

22 Yen, S. T. et al. Somatic mosaicism and allele complexity induced by CRISPR/Cas9 RNA injections in mouse zygotes. Dev Biol 393, 3-9, doi:10.1016/j.ydbio.2014.06.017 (2014).

23 Girard, J., Ferre, P., Pegorier, J. P. & Duee, P. H. Adaptations of glucose and fatty acid metabolism during perinatal period and suckling-weaning transition. Physiol Rev 72, 507-562, doi:10.1152/physrev.1992.72.2.507 (1992).

24 Hillman, N. H., Kallapur, S. G. & Jobe, A. H. Physiology of transition from intrauterine to extrauterine life. Clin Perinatol 39, 769-783, doi:10.1016/j.clp.2012.09.009 (2012).

25 Wang, N. D. et al. Impaired energy homeostasis in C/EBP alpha knockout mice. Science 269, 1108-1112 (1995).

26 Ibdah, J. A. et al. Lack of mitochondrial trifunctional protein in mice causes neonatal hypoglycemia and sudden death. J Clin Invest 107, 1403-1409, doi:10.1172/JCI12590 (2001).

27 Cotter, D. G., ďAvignon, D. A., Wentz, A. E., Weber, M. L. & Crawford, P. A. Obligate role for ketone body oxidation in neonatal metabolic homeostasis. J Biol Chem 286, 6902-6910, doi:10.1074/jbc.M110.192369 (2011).

28 Vafai, S. B. & Mootha, V. K. Mitochondrial disorders as windows into an ancient organelle. Nature 491, 374-383, doi:10.1038/nature11707 (2012).

29 Rinaldo, P., Matern, D. & Bennett, M. J. Fatty acid oxidation disorders. Annu Rev Physiol 64, 477-502, doi:10.1146/annurev.physiol.64.082201.154705 (2002).

30 Schuler, A. M. & Wood, P. A. Mouse models for disorders of mitochondrial fatty acid beta-oxidation. ILAR J 43, 57-65 (2002).

31 Maaswinkel-Mooij, P. D. et al. Depletion of mitochondrial DNA in the liver of a patient with lactic acidemia and hypoketotic hypoglycemia. J Pediatr 128, 679-683 (1996).

32 Przyrembel, H. et al. Glutaric aciduria type II: report on a previously undescribed metabolic disorder. Clin Chim Acta 66, 227-239 (1976).

33 Vockley, J., Rinaldo, P., Bennett, M. J., Matern, D. & Vladutiu, G. D. Synergistic heterozygosity: disease resulting from multiple partial defects in one or more metabolic pathways. Mol Genet Metab 71, 10-18, doi:10.1006/mgme.2000.3066 (2000).

34 Ney, P. A. Mitochondrial autophagy: Origins, significance, and role of BNIP3 and NIX. Biochim Biophys Acta 1853, 2775-2783, doi:10.1016/j.bbamcr.2015.02.022 (2015).

35 Geissler, A. et al. The mitochondrial presequence translocase: an essential role of Tim50 in directing preproteins to the import channel. Cell 111, 507-518 (2002).

36 Dickinson, M. E. et al. High-throughput discovery of novel developmental phenotypes. Nature 537, 508-514, doi:10.1038/nature19356 (2016).

37 Reyes, A. et al. Mutations in TIMM50 compromise cell survival in OxPhos-dependent metabolic conditions. EMBO Mol Med, doi:10.15252/emmm.201708698 (2018).

38 Yamamoto, H. et al. Tim50 is a subunit of the TIM23 complex that links protein translocation across the outer and inner mitochondrial membranes. Cell 111, 519-528 (2002).

39 Ting, S. Y., Yan, N. L., Schilke, B. A. & Craig, E. A. Dual interaction of scaffold protein Tim44 of mitochondrial import motor with channel-forming translocase subunit Tim23. Elife 6, doi:10.7554/eLife.23690 (2017).

40 Weidberg, H. & Amon, A. MitoCPR-A surveillance pathway that protects mitochondria in response to protein import stress. Science 360, doi:10.1126/science.aan4146 (2018).

41 Wiedemann, N., Frazier, A. E. & Pfanner, N. The protein import machinery of mitochondria. J Biol Chem 279, 14473-14476, doi:10.1074/jbc.R400003200 (2004).

42 Calvo, S. E. et al. Comparative Analysis of Mitochondrial N-Termini from Mouse, Human, and Yeast. Mol Cell Proteomics 16, 512-523, doi:10.1074/mcp.M116.063818 (2017).

43 Rhee, H. W. et al. Proteomic mapping of mitochondria in living cells via spatially restricted enzymatic tagging. Science 339, 1328-1331, doi:10.1126/science.1230593 (2013).

44 Baumgartner M.R., V. D., and C. Dionisi-Vici. in Inborn Metabolic Diseases (ed J.-M. Saudubray) Ch. 21, 321-331 (Springer-Verlag, 2016).

45 El-Hattab, A. W. Inborn errors of metabolism. Clin Perinatol 42, 413-439, x, doi:10.1016/j.clp.2015.02.010 (2015).

46 Kanehisa, M., Sato, Y., Kawashima, M., Furumichi, M. & Tanabe, M. KEGG as a reference resource for gene and protein annotation. Nucleic Acids Res 44, D457-462, doi:10.1093/nar/gkv1070 (2016).

47 Oza, J. P. et al. Robust production of recombinant phosphoproteins using cell-free protein synthesis. Nat Commun 6, 8168, doi:10.1038/ncomms9168 (2015).

48 Pirman, N. L. et al. A flexible codon in genomically recoded Escherichia coli permits programmable protein phosphorylation. Nat Commun 6, 8130, doi:10.1038/ncomms9130 (2015).

49 Stojanovski, D., Pfanner, N. & Wiedemann, N. Import of proteins into mitochondria. Methods Cell Biol 80, 783-806, doi:10.1016/S0091-679X(06)80036-1(2007).

50 Murakami, H., Pain, D. & Blobel, G. 70-kD heat shock-related protein is one of at least two distinct cytosolic factors stimulating protein import into mitochondria. J Cell Biol 107, 2051-2057 (1988).

51 Hachiya, N. et al. MSF, a novel cytoplasmic chaperone which functions in precursor targeting to mitochondria. EMBO J 13, 5146-5154 (1994).

52 Mossmann, D. et al. Amyloid-beta peptide induces mitochondrial dysfunction by inhibition of preprotein maturation. Cell Metab 20, 662-669, doi:10.1016/j.cmet.2014.07.024 (2014).

53 Mukhopadhyay, A., Yang, C. S., Wei, B. & Weiner, H. Precursor protein is readily degraded in mitochondrial matrix space if the leader is not processed by mitochondrial processing peptidase. J Biol Chem 282, 37266-37275, doi:10.1074/jbc.M706594200 (2007).

54 Schatton, D. et al. CLUH regulates mitochondrial metabolism by controlling translation and decay of target mRNAs. J Cell Biol 216, 675-693, doi:10.1083/jcb.201607019 (2017).

55 Schmidt, O. et al. Regulation of mitochondrial protein import by cytosolic kinases. Cell 144, 227-239, doi:10.1016/j.cell.2010.12.015 (2011).

56 Rao, S. et al. Biogenesis of the preprotein translocase of the outer mitochondrial membrane: protein kinase A phosphorylates the precursor of Tom40 and impairs its import. Mol Biol Cell 23, 1618-1627, doi:10.1091/mbc.E11-11-0933 (2012).

57 Gerbeth, C. et al. Glucose-induced regulation of protein import receptor Tom22 by cytosolic and mitochondria-bound kinases. Cell Metab 18, 578-587, doi:10.1016/j.cmet.2013.09.006 (2013).

58 Harbauer, A. B. et al. Mitochondria. Cell cycle-dependent regulation of mitochondrial preprotein translocase. Science 346, 1109-1113, doi:10.1126/science.1261253 (2014).

59 Acin-Perez, R. et al. Cyclic AMP produced inside mitochondria regulates oxidative phosphorylation. Cell Metab 9, 265-276, doi:10.1016/j.cmet.2009.01.012 (2009).

60 Fang, J. K. et al. Site specific phosphorylation of cytochrome c oxidase subunits I, IVi1 and Vb in rabbit hearts subjected to ischemia/reperfusion. FEBS Lett 581, 1302-1310, doi:10.1016/j.febslet.2007.02.042 (2007).

61 Lefkimmiatis, K., Leronni, D. & Hofer, A. M. The inner and outer compartments of mitochondria are sites of distinct cAMP/PKA signaling dynamics. J Cell Biol 202, 453-462, doi:10.1083/jcb.201303159 (2013).

62 May, T. & Soll, J. 14-3-3 proteins form a guidance complex with chloroplast precursor proteins in plants. Plant Cell 12, 53-64 (2000).

63 Waegemann, K. & Soll, J. Phosphorylation of the transit sequence of chloroplast precursor proteins. J Biol Chem 271, 6545-6554 (1996).

64 Robin, M. A., Prabu, S. K., Raza, H., Anandatheerthavarada, H. K. & Avadhani, N. G. Phosphorylation enhances mitochondrial targeting of GSTA4-4 through increased affinity for binding to cytoplasmic Hsp70. J Biol Chem 278, 18960-18970, doi:10.1074/jbc.M301807200 (2003).

65 Anandatheerthavarada, H. K., Sepuri, N. B. & Avadhani, N. G. Mitochondrial targeting of cytochrome P450 proteins containing NH2-terminal chimeric signals involves an unusual TOM20/TOM22 bypass mechanism. J Biol Chem 284, 17352-17363, doi:10.1074/jbc.M109.007492 (2009).

66 Lee, J., O’Neill, R. C., Park, M. W., Gravel, M. & Braun, P. E. Mitochondrial localization of CNP2 is regulated by phosphorylation of the N-terminal targeting signal by PKC: implications of a mitochondrial function for CNP2 in glial and non-glial cells. Mol Cell Neurosci 31, 446-462, doi:10.1016/j.mcn.2005.10.017 (2006).

67 Zhao, X. et al. Phosphoproteome analysis of functional mitochondria isolated from resting human muscle reveals extensive phosphorylation of inner membrane protein complexes and enzymes. Mol Cell Proteomics 10, M110 000299, doi:10.1074/mcp.M110.000299 (2011).

68 Chung, J. et al. Erythropoietin signaling regulates heme biosynthesis. Elife 6, doi:10.7554/eLife.24767 (2017).

69 De Rasmo, D., Panelli, D., Sardanelli, A. M. & Papa, S. cAMP-dependent protein kinase regulates the mitochondrial import of the nuclear encoded NDUFS4 subunit of complex I. Cell Signal 20, 989-997, doi:10.1016/j.cellsig.2008.01.017 (2008).

70 Wang, L. et al. Cloning and mitochondrial localization of full-length D-AKAP2, a protein kinase A anchoring protein. Proc Natl Acad Sci U S A 98, 3220-3225, doi:10.1073/pnas.051633398 (2001).

71 Alam, R. et al. cDNA cloning and characterization of mitochondrial import stimulation factor (MSF) purified from rat liver cytosol. J Biochem 116, 416-425 (1994).

72 Komiya, T., Hachiya, N., Sakaguchi, M., Omura, T. & Mihara, K. Recognition of mitochondria-targeting signals by a cytosolic import stimulation factor, MSF. J Biol Chem 269, 30893-30897 (1994).

73 Johnson, C. et al. Visualization and biochemical analyses of the emerging mammalian 14-3-3-phosphoproteome. Mol Cell Proteomics 10, M110 005751, doi:10.1074/mcp.M110.005751 (2011).

74 La, P., Yang, G. & Dennery, P. A. Mammalian target of rapamycin complex 1 (mTORC1)-mediated phosphorylation stabilizes ISCU protein: implications for iron metabolism. J Biol Chem 288, 12901-12909, doi:10.1074/jbc.M112.424499 (2013).

75 Horn, H. et al. KinomeXplorer: an integrated platform for kinome biology studies. Nat Methods 11, 603-604, doi:10.1038/nmeth.2968 (2014).

76 Kerk, D., Silver, D., Uhrig, R. G. & Moorhead, G. B. “PP2C7s”, Genes Most Highly Elaborated in Photosynthetic Organisms, Reveal the Bacterial Origin and Stepwise Evolution of PPM/PP2C Protein Phosphatases. PLoS One 10, e0132863, doi:10.1371/journal.pone.0132863 (2015).

77 Chen, M. J., Dixon, J. E. & Manning, G. Genomics and evolution of protein phosphatases. Sci Signal 10, doi:10.1126/scisignal.aag1796 (2017).

78 Duncan, M. R., Fullerton, M. & Chaudhuri, M. Tim50 in Trypanosoma brucei possesses a dual specificity phosphatase activity and is critical for mitochondrial protein import. J Biol Chem 288, 3184-3197, doi:10.1074/jbc.M112.436378 (2013).

79 Lombard, D. B. et al. Mammalian Sir2 homolog SIRT3 regulates global mitochondrial lysine acetylation. Mol Cell Biol 27, 8807-8814, doi:10.1128/MCB.01636-07 (2007).

80 Yu, J. et al. Metabolic characterization of a Sirt5 deficient mouse model. Sci Rep 3, 2806, doi:10.1038/srep02806 (2013).

81 Veling, M. T. et al. Multi-omic Mitoprotease Profiling Defines a Role for Oct1p in Coenzyme Q Production. Mol Cell 68, 970-977 e911, doi:10.1016/j.molcel.2017.11.023 (2017).

82 Stefely, J. A. et al. Mitochondrial protein functions elucidated by multi-omic mass spectrometry profiling. Nat Biotechnol 34, 1191-1197, doi:10.1038/nbt.3683 (2016).

83 Linstrom, J. NIST Standard Reference Database Number 69. (2014).

84 Hutchins, P. D., Russell, J. D. & Coon, J. J. LipiDex: An Integrated Software Package for High-Confidence Lipid Identification. Cell Syst 6, 621-625 e625, doi:10.1016/j.cels.2018.03.011 (2018).

85 Fernandez, M., Mano, S., de Fernando, D. G., Ordonez, J. A. & Hoz, L. Use of beta-hydroxyacyl-CoA-dehydrogenase (HADH) activity to differentiate frozen from unfrozen fish and shellfish. Eur Food Res Technol 209, 205-208, doi:DOI 10.1007/s002170050481 (1999).

86 Hebert, A. S. et al. The one hour yeast proteome. Mol Cell Proteomics 13, 339-347, doi:10.1074/mcp.M113.034769 (2014).

87 Riley, N. M. et al. Phosphoproteomics with Activated Ion Electron Transfer Dissociation. Anal Chem 89, 6367-6376, doi:10.1021/acs.analchem.7b00212 (2017).

88 Geer, L. Y. et al. Open mass spectrometry search algorithm. J Proteome Res 3, 958-964, doi:10.1021/pr0499491 (2004).

89 Wenger, C. D., Phanstiel, D. H., Lee, M. V., Bailey, D. J. & Coon, J. J. COMPASS: a suite of pre-and post-search proteomics software tools for OMSSA. Proteomics 11, 1064-1074, doi:10.1002/pmic.201000616 (2011).

90 Taus, T. et al. Universal and confident phosphorylation site localization using phosphoRS. J Proteome Res 10, 5354-5362, doi:10.1021/pr200611n (2011).

91 Meisinger, C., Pfanner, N. & Truscott, K. N. Isolation of yeast mitochondria. Methods Mol Biol 313, 33-39, doi:10.1385/1-59259-958-3:033 (2006).

92 Johnston, A. J. et al. Insertion and assembly of human tom7 into the preprotein translocase complex of the outer mitochondrial membrane. J Biol Chem 277, 42197-42204, doi:10.1074/jbc.M205613200 (2002).

93 Kwon, Y. C. & Jewett, M. C. High-throughput preparation methods of crude extract for robust cell-free protein synthesis. Sci Rep 5, 8663, doi:10.1038/srep08663 (2015).

94 Caschera, F. & Noireaux, V. Preparation of amino acid mixtures for cell-free expression systems. Biotechniques 58, 40-43, doi:10.2144/000114249 (2015).

95 Floyd, B. J. et al. Mitochondrial Protein Interaction Mapping Identifies Regulators of Respiratory Chain Function. Mol Cell 63, 621-632, doi:10.1016/j.molcel.2016.06.033 (2016).

